# Lineage origin of spinal cord cell type diversity

**DOI:** 10.64898/2026.02.12.705305

**Authors:** Sophie A. Gobeil, Francisco Da Silveira Neto, Giulia Silvestrelli, Matthijs Smits, Carmen Streicher, Giselle Cheung, Simon Hippenmeyer, Lora B. Sweeney

## Abstract

The complexity and specificity of movement in vertebrates is driven by a rich diversity of spinal motor and interneuron cell types. During development, eleven spinal cord progenitor domains generate an equivalent number of cardinal neuron types. How progenitor domains, individual progenitors, and post-mitotic diversity relate is still unknown. We performed high-resolution, single-progenitor cell lineage tracing in the embryonic mouse spinal cord using mosaic analysis with double markers (MADM). Our quantitative study of lineage progression revealed that spinal cord progenitors undergo highly variable numbers of proliferative, neurogenic, and gliogenic cell divisions. The nascent clonally-related neurons migrate radially over large distances, span the dorsoventral axis, and even cross the midline, demonstrating striking bilaterality. Molecular and morphometric analysis indicate high levels of progenitor multipotency, with an individual progenitor capable of producing several molecularly and morphologically distinct neuron types, as well as astrocytes. These findings redefine spinal cord development as a process in which lineage variability—rather than rigid progenitor identity—drives the generation of cellular diversity.

## INTRODUCTION

Developing organisms face many challenges in building a nervous system: consistently producing the correct types of neurons and glia, balancing them in the right proportions, organizing their migration, and forming synapses between the appropriate partners. Across species and neural tissues, various developmental strategies are employed to accomplish these tasks. In invertebrates, exemplified by the *Drosophila* brain and ventral nerve cord, neuronal identity and function are tightly regulated by deterministic neuroblast lineages governed by temporal transcriptional programs and spatial position^1,2^. In contrast, in the vertebrate retina, multipotent neural progenitors can produce every retinal neuron type, with birth order, extrinsic, and stochastic factors influencing the composition and number of progeny^3–7^.

Remarkably, despite advances in understanding lineage progression in the developing brain and retina, the role of lineage in generating cell-type diversity in the spinal cord is still unknown. How lineage links spinal progenitor and neuron types, how progenitors generate variant subtypes of neurons and glia, and to what extent lineage relationships constrain cell fate and regulate consistent cell-type proportions are still unanswered.

These questions are particularly relevant to the spinal cord, due to its rich diversity of cell types comprising numerous molecularly distinct spinal interneurons (INs) and motor neurons (MNs) which receive sensory information and coordinate motor output. Such extensive molecular diversity seems to be a defining feature of spinal neurons, underlying their functional specialization^8,9^. They are commonly classified into cardinal classes based on their spatial origin in the embryonic neural tube^10–13^. During development, this cardinal architecture is driven by opposing morphogen gradients, which pattern the dorsoventral axis of the neural tube into eleven progenitor domains that each generate a population of neurons. Within each of these domains, temporal transcriptional programs serve as an additional axis driving neuron diversification^14–16^. Despite such well-established early developmental drivers and extensive post-mitotic neural diversity however, the precise mechanism by which individual progenitors generate such diversity still remains poorly understood.

Classical loss- and gain-of-function studies and fate-mapping approaches have provided population-level insight into spinal progenitor potential, but lack the resolution to reconstruct the full developmental trajectory of individual cells^17^. Early retroviral lineage tracing work in chick demonstrated that MN progenitors can produce both MNs and INs^18–20^. More recent genetic labeling further showed that neural progenitors can switch identities in response to dynamic morphogen signaling, blurring the boundaries between domains^21–23^. However, these studies were limited by either a lack of molecular resolution, restriction to certain classes, or debate about true clonality. Fundamentally, for the spinal cord, the lineage relationship between each of the eleven progenitor types, the eleven cardinal neuron classes, and the diversity of subpopulations within each class is still unclear.

The development of new methodologies for single-progenitor lineage tracing have begun to parse the relative contribution of lineage to the diversity of cell types in the mammalian brain, setting the stage for parallel analysis in the spinal cord. Mosaic analysis with double markers (MADM)^24,25^ has yielded high-precision, single-progenitor lineage traces of neuro- and gliogenic dynamics. Its quantitative precision, color-coding the two descendent hemilineages of a dividing progenitor cell, finds contrasting developmental strategies by brain region: exemplified by the highly stereotyped lineage progression of the cerebral cortex^26^ and the more variable pattern of cell-type generation in the superior colliculus^27^.

Here, we aim to uncover the principles of lineage progression and cell-type diversification in the mouse spinal cord using MADM. Capitalizing on the single-cell and unambiguous clonal resolution of MADM, we dissected the full lineage output and spatial organization of individual spinal neural progenitors spanning the dorsoventral and rostrocaudal axes. Subsequent high-resolution, volumetric images of intact spinal cords further resolved three-dimensional spatial and morphological information of clones *in situ*, providing a basis to relate lineage, spinal circuit organization, and function. We observed striking variability in progenitor division behavior and clonal dispersion: (i) individual progenitors generated diverse neuronal and glial types; (ii) clonally-related neurons spanned the dorsoventral axis; and (iii) neurons derived from one clone even crossed the midline. Lineage variability and multipotency are thus unexpectedly central drivers of cellular diversity during spinal cord development.

## RESULTS

### Individual multipotent progenitors produce neurons and glia in the developing spinal cord

To resolve the origins of spinal neurons and glia and elucidate the developmental strategy for generating cell diversity in the spinal cord (**Fig. 1a**), we used mosaic analysis with double markers (MADM)^24,25^, a means to visualize the lineages of individual neural progenitors (NPs) in the mouse embryonic neural tube (**Fig. 1b**). MADM relies on Cre-mediated recombination of a chimeric reporter cassette in dividing progenitor cells (**Fig. S1a**), resulting in the expression of either a red (tdT) or green (GFP) fluorescent reporter in each daughter hemilineage. *Sox2-CreER,* universally expressed in NPs^28^, was used as a driver to induce MADM labeling at embryonic day 9.5 (E9.5), E10.5, and E11.5, spanning early to mid spinal neurogenesis (**Fig. S1b**). Spinal cords were collected at E18.5, when neurogenesis is complete and the nascent neurons have reached their final settling positions^29–31^ and assumed their mature multipolar morphologies^32^. Spinal cord samples were cleared and imaged in wholemount, exploiting the high resolution and penetration depth of lightsheet imaging to resolve MADM clones in their entirety.

**Figure 1.**
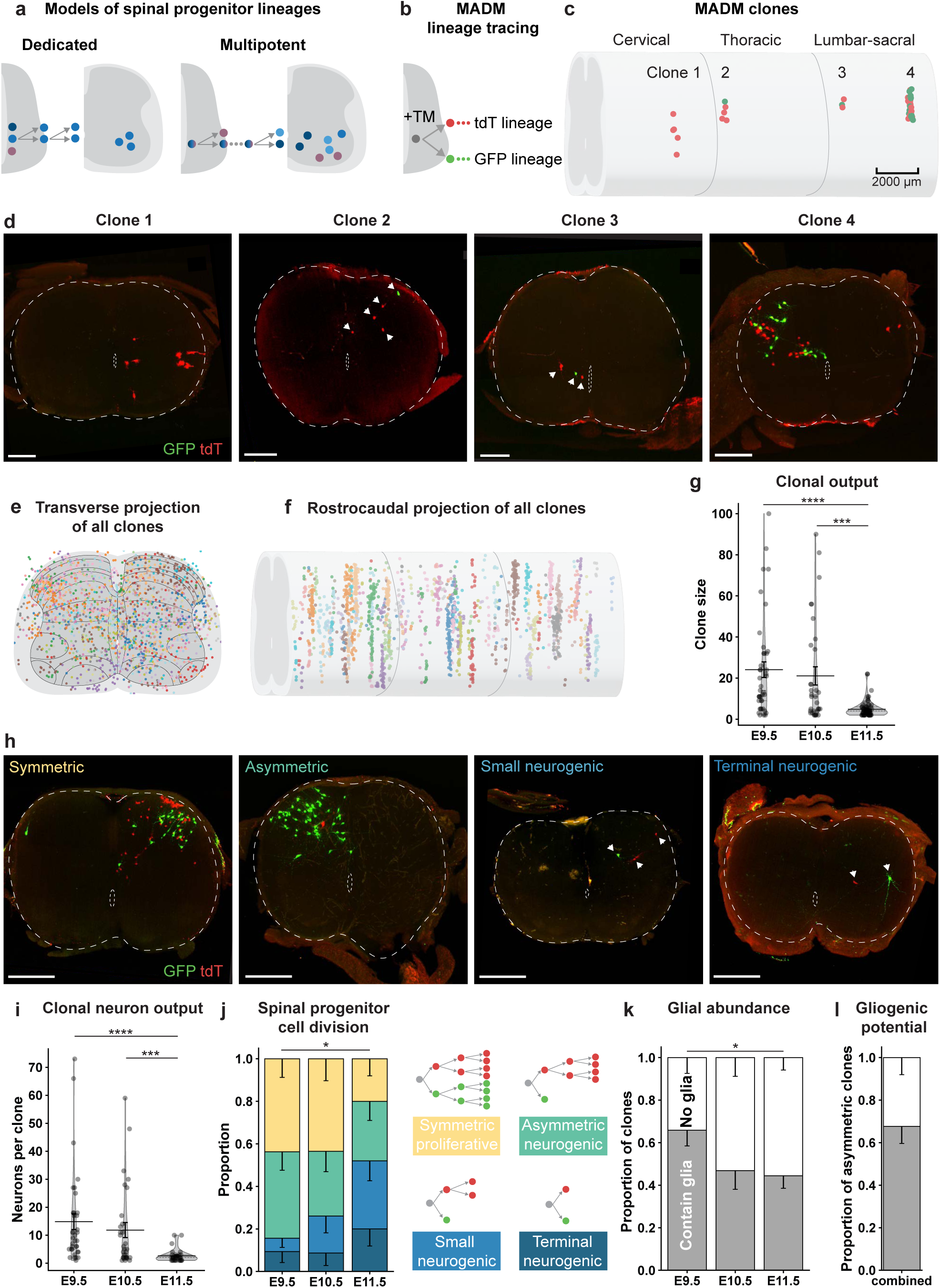
MADM-based clonal analysis reveals multipotent spinal cord progenitors produce neurons and glia. **a)** Putative models of spinal progenitor lineage progression via dedicated versus multipotent progenitors. **b)** Simplified schematic of MADM lineage tracing. **c)** Representative spinal cord with four MADM-labeled clones (red/green), induced at E9.5. **d)** Maximum intensity projections of the clones in (**c**) with tdT and GFP hemilineages. From left to right: cervical level; 250 μm projection, thoracic level; 200 μm projection, lumbar level; 200 μm projection, sacral level; 300 μm projection. Scale bars, 200 μm. **e)** Transverse projections of all clones across E9.5, E10.5, and E11.5 induction time points. Each colour depicts an individual clone. *n* = 145. **f)** Longitudinal projections of all clones. **g)** Size of clones (total number of cells) across induction time points. E9.5 (*n* = 41), E10.5 (*n* = 32), E11.5 (*n* = 72). Kruskal-Wallis test with Dunn’s multiple comparisons test; *p* < 0.0001; *p* = 0.4404 between E9.5 and E10.5; p < 0.0001 between E9.5 and E11.5; *p* = 0.0003 between E10.5 and E11.5. **h)** Example maximum intensity projections of MADM clones reflecting symmetric (TM E10.5, lumbar level; 300 μm projection), asymmetric (TM E9.5, sacral level; 350 μm projection), small neurogenic (TM E10.5, cervical level; 150 μm projection), and terminal neurogenic (TM E9.5, lumbar level; 300 μm projection) division patterns. tdT and GFP hemilineages are labeled. Scale bars, 200 μm. **i)** Clonal neuron output (number of neurons per neuron-containing clone) across induction time points. E9.5 (*n* = 35), E10.5 (*n* = 30), E11.5 (*n* = 30). Kruskal-Wallis test with Dunn’s multiple comparisons test; *p* < 0.0001; *p* = 0.3954 between E9.5 and E10.5; *p* < 0.0001 between E9.5 and E11.5; *p* = 0.0007 between E10.5 and E11.5. **j)** Proportion of neurogenic clones with symmetric (≥4 G; ≥4 R), asymmetric (≤3 G; ≥4 R), small neurogenic (≤4 G; ≤4 R), and terminal neurogenic (1 G; 1 R) division patterns across induction time points. GFP and tdT hemilineages are interchangeable in division pattern definitions. E9.5 (*n* = 32), E10.5 (*n* = 23), E11.5 (*n* = 25). Fisher’s exact test; *p* = 0.6072 between E9.5 and E10.5; *p* = 0.2673 between E10.5 and E11.5; *p* = 0.0236 between E9.5 and E11.5. **k)** Proportion of all clones containing or not containing glia across induction time points. E9.5 (*n* = 41), E10.5 (*n* = 32), E11.5 (*n* = 72). Fisher’s exact test; *p* = 0.1521 between E9.5 and E10.5; *p* = 0.8341 between E10.5 and E11.5; *p* = 0.0327 between E9.5 and E11.5. **l)** Proportion of asymmetric clones containing or not containing glia. *n* = 44. Violin plot data are presented as mean ± SEM. Proportions are presented with SE of the proportion (lower limit). All clones were analyzed at E18.5. *n* indicates the number of clones. \**p* < 0.05; \*\*\**p* < 0.001; \*\*\*\**p* < 0.0001. **Abbreviations:** TM, tamoxifen; SE, standard error; SEM, standard error of the mean; E, embryonic day; GFP, green fluorescent protein; tdT, tdTomato; MADM, mosaic analysis with double markers.

Discrete clusters of red/green (R/G)-positive cells were sparsely distributed along the length of the spinal cord (**Fig. 1c-d; S1c**). Stringent criteria were used to identify clones (**Fig. S1d-j**; see **Clonal Resolution** in **Methods**). In total, 145 R/G clones were detected in ∼50 animals that tiled the spinal cord’s transverse and rostrocaudal axes (**Fig. 1e,f**). While transverse spread of clonally related cells could be extensive, rostrocaudal dispersion was restricted (**Fig. S1f-g**). We focused our analysis on these R/G clones because they reveal the division patterns of labeled NPs and provided insight into their cellular output potential. Clone size, the total cells produced by a single NP, decreased from E9.5 to E11.5, reflecting a decrease in progenitor potential over time (**Fig. 1g**), consistent with what has been observed in the brain^26,27^. Despite this reduction however, total cell and neuron output in the spinal cord was highly variable (**Fig. 1g,i; S2a**), differing from the stereotypy of cortical lineage output^26^.

MADM R/G hemilineage color-coding enabled us to quantify progenitor cell division patterns in detail. We classified clone architectures into four patterns: *symmetric clones* with two large hemilineages reflecting self-renewing progenitor divisions; *asymmetric neurogenic clones* in which the minority hemilineage contained one to three, and the majority hemilineage contained many, cells; *small neurogenic clones* with three or more cells and at least one neuron; and *terminal neurogenic* two-cell clones of one to two neurons (**Fig. 1h,j**). From E9.5 to E11.5, we observed a transition toward more neurogenic progenitor division patterns (**Fig. 1j**). However, at all induction timepoints, progenitors were engaged in every mode of progenitor cell division. The size of symmetric and asymmetric clones, which reflect the number of rounds of proliferative and neurogenic divisions, respectively, were also variable (**Fig. S2b-c**), a striking difference from the consistency of cell division patterns in other brain regions such as the cortex^26^. In summary, the spinal progenitor population behaves heterogeneously, undergoing variable rounds of proliferative and neurogenic divisions.

In addition to neurons, MADM clones also frequently contained other spinal cell types, including glia, septal cells, and progenitors, identified based on their morphology (**Fig. S3a-d**). The majority of clones contained glia (∼45-65%; **Fig. 1k**). Among asymmetric neurogenic clones, an exceptionally high proportion of ∼70% contained glia, indicating that a majority of progenitors proceed to gliogenesis following neurogenesis (**Fig. 1l**). This contrasts with the cortex and superior colliculus in which just under 20% and 15% switch from neuro- to gliogenesis, respectively^26,27^. Glia-only clones comprised only ∼30% of clones, and were observed only after E11.5 induction (**Fig. S3c**). Glia output, like neuron output, was also strikingly variable (**Fig. S4a-b**). Glia-containing clones often contained multiple types of glia and astrocytes (**Fig. S4c**). The majority of spinal progenitors are thus multipotent, producing both neurons and several types of glia.

### Spinal neurons and glia settle radially and bilaterally

The clonal resolution achieved with MADM made it possible to investigate how clone architecture contributes to the developmental and cellular organization of the spinal cord. How do clonally-related cells disperse and settle in the developing neural tube? Do clonal units adopt a consistent structure? To begin to answer these questions, we first extended our analysis to score the settling position of cells within a clone. Initial spatial analysis revealed clones of different sizes and cell-type composition that adopted a radial arrangement, spanning the distance between the central canal and pia (**Fig. 2b**). However, clonally related cells were also found to spread widely in the transverse plane and had varied architecture **(Fig. 2b; Fig. S1f**). Thus, although clonally related cells often dispersed radially, there were no strict constraints on clone structure, as observed in the cortex^26^.

**Figure 2.**
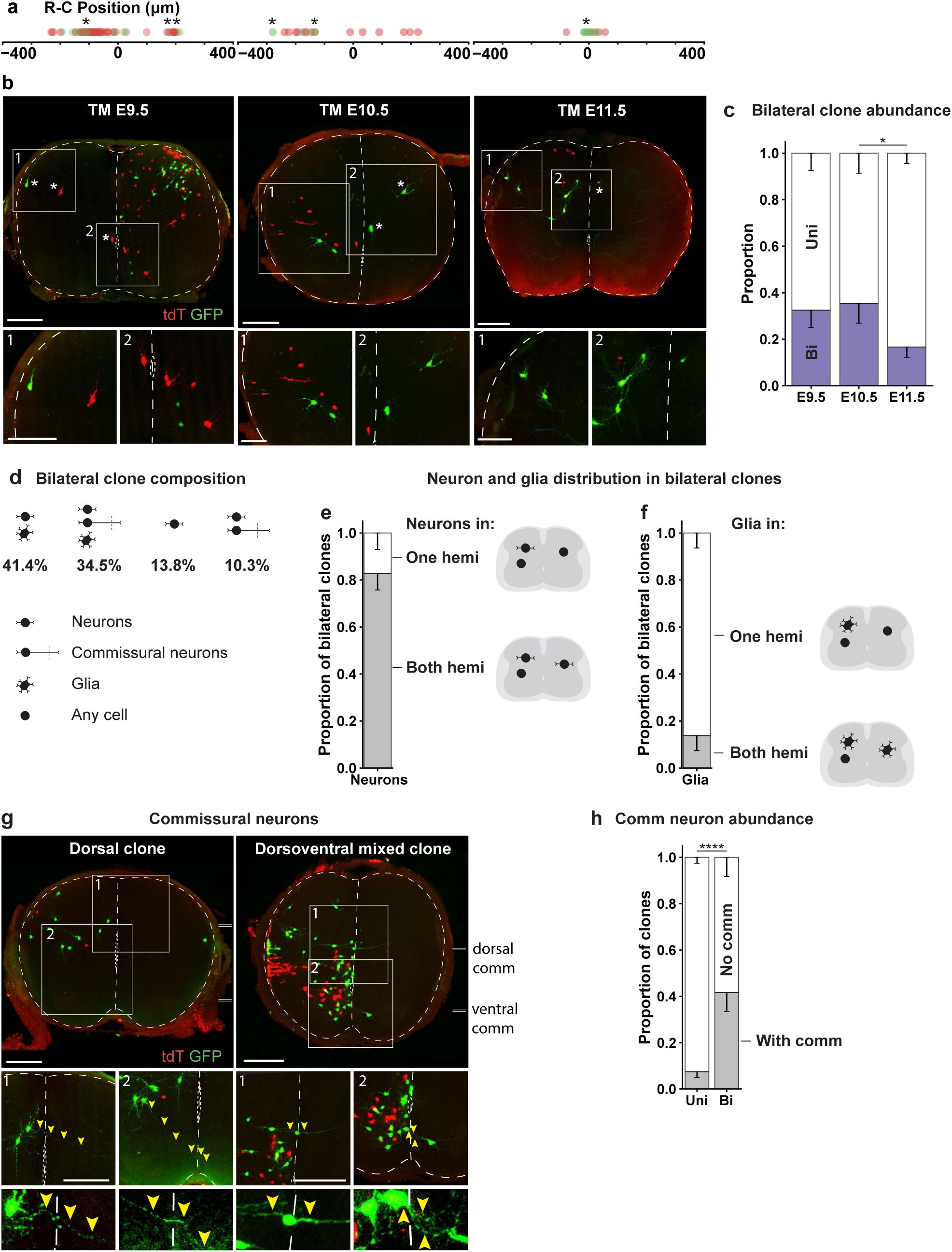
MADM clones show radial and bilateral clone architecture. **a)** Distribution of clonally related cells in (**b**) along the rostrocaudal axis. Each dot represents a single cell within the clone. Asterisks represent selected cells also marked in (**b**). **b)** Maximum intensity projections of bilateral MADM clones induced at E9.5 (lumbar level; overview 400 μm projection; insets 200 μm projection), E10.5 (thoracic level; 150 μm projection), and E11.5 (lumbar level; 150 μm projection). tdT and GFP hemilineages are labeled. Of note, the TM E11.5 is the sole clone in a single-clone spinal cord. **c)** Proportions of unilateral and bilateral clones across labeling time points. TM E9.5 (*n* = 40), E10.5 (*n* = 31), E11.5 (*n* = 72). Fisher’s exact test; *p* = 0.8058 between E9.5 and E10.5; *p* = 0.0427 between E10.5 and E11.5; *p* = 0.0621 between E9.5 and E11.5. **d)** Proportion of bilateral clones with different neuron, commissural neuron, and glial compositions. Cells of uncertain identity excluded from quantification. Bilateral clones in which a hemisphere contains only cells of uncertain identity excluded from cell-type quantifications. *n* = 29. Icons corresponding to neurons, commissural neurons, glia, or any cell type are defined in the figure. **e)** Proportion of neuron-containing bilateral clones comprising neurons in one hemisphere (hemi.) or both hemispheres. *n* = 29. Icons depicting cell-types are defined in (**d**). **f)** Proportion of glia-containing bilateral clones comprising glia in one or both hemispheres. *n* = 22. Icons depicting cell-types are defined in (**d**). **g)** Example maximum intensity projections of bilateral clones containing commissural neurons. Dorsal clone: TM E10.5, thoracic level; overview 200 μm projection; insets 200 μm projection. Dorsoventral mixed clone: TM E9.5, thoracic level; overview 200 μm projection; insets 75 and 150 μm projections. tdT and GFP hemilineages are labeled. The level of the dorsal and ventral commissures are indicated by double lines. **h)** Proportions of unilateral and bilateral clones containing commissural neurons. Unilateral (*n* = 107), bilateral (*n* = 36). Fisher’s exact test; *p* < 0.0001. Violin plot data are presented as mean ± SEM. Proportions are presented with SE of the proportion (lower limit). All clones were analyzed at E18.5. *n* indicates the number of clones. \**p* < 0.05; \*\*\*\**p* < 0.0001. **Abbreviations:** TM, tamoxifen; SE, standard error; SEM, standard error of the mean; E, embryonic day; GFP, green fluorescent protein; tdT, tdTomato; Bi, bilateral; Uni, unilateral; hemi, hemisphere; comm, commissural; R-C, rostrocaudal; MADM, mosaic analysis with double markers.

Strikingly, although most clones were restricted to a single spinal cord hemisphere, many were bilateral (**Fig. 2c**). These contained cells that settled in both hemispheres at a similar rostrocaudal level (**Fig. 2a-b**). Contralateral cells were often similar in dorsoventral level to those on the ipsilateral side (**Fig. 2b; S5a**). Bilateral clones ranged in size, but were on average larger than unilateral clones (**Fig. S5b**). In large bilateral clones, cell numbers were often unevenly distributed between the two hemispheres (**Fig. S5c**). Bilateral clones typically occupied dorsal regions or broadly spanned the DV axis, and were rarely ventrally restricted (**Fig. 2b,g; S5d**). Their proportions were consistent across rostrocaudal levels (**Fig. S5e**). They frequently contained both glia and neurons (**Fig. 2d-f**), and were enriched in commissural neurons (**Fig. 2g-h**). No clear hemisphere-specific segregation by cell type (neuron, commissural neuron, or glial) was observed (**Fig. S5f**). And typically, cells of both hemilineages were present in one hemisphere but only of one hemilineage in the other (**Fig. S5g**). Bilateral clones were sometimes the sole clone in single-clone-containing spinal cords, supporting that they arise from a single recombination event rather than two adjacent events (**Fig. 2b; S5a**). Together, these observations indicate that individual neural progenitors can generate broadly dispersing bilateral lineages with varied settling patterns. Such extensive bilaterality has not been previously reported in the spinal cord or in clonal analysis of the cortex^26^ or superior colliculus^27^.

### Clonally related neurons display diverse morphologies

The high variability and extent of clonal dispersion argues against a simple relationship between clone architecture and particular cell-types in the spinal cord. We therefore asked whether progenitor lineage instead constrains other functionally relevant neuron features, such as morphology^33–35^. We reconstructed neurons from MADM-labeled clones and studied their proximal morphology, quantifying neurite geometry, number of primary neurites, and branching complexity within a 100 μm radius (**Fig. 3a-b; S6a-b**). Based on these features, neurons were classified into distinct morphotypes that capture key differences in structure^36–38^ (**Fig. 3b; S6d-h**). Following these criteria, we successfully identified motor neurons in our dataset as the complex stellate morphotype, confirming a link between our described morphotypes and established molecular/functional spinal neuron identities (**Fig. S5c**).

**Figure 3.**
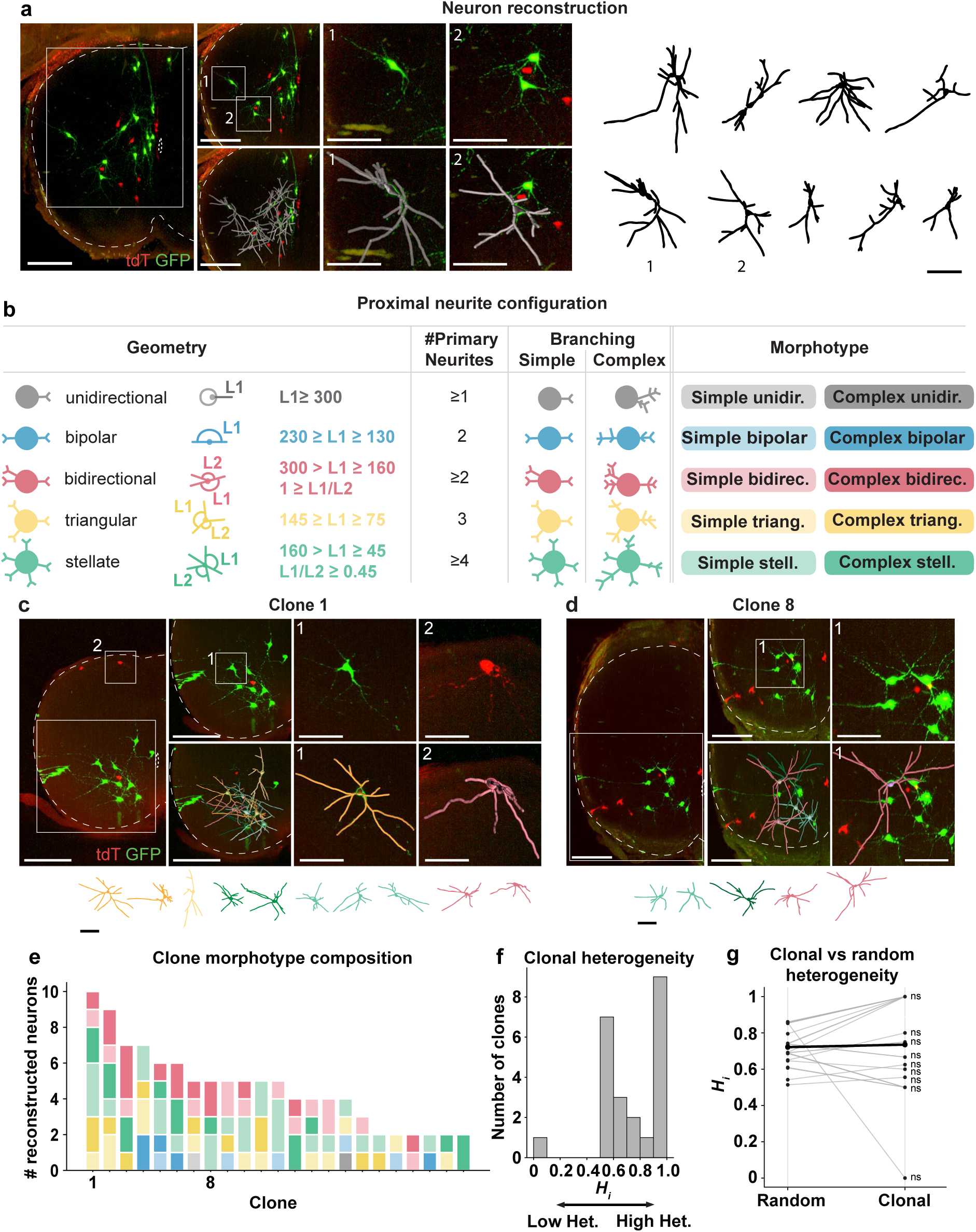
Morphological divergence of clonally related neurons. **a)** Maximum intensity projection of a MADM clone induced at E9.5 (thoracic level; 300 μm projection). Nine of 25 labeled neurons (36%) were reconstructed. tdT and GFP hemilineages are labeled. Scale bars, 200 μm (image), 100 μm (traces). **b)** Classification criteria for spinal neuron morphotypes. L1: largest angle between neighboring primary neurites; L2: second largest angle between neighboring primary neurites. Adapted from Fukuda et al., 2020. **c-d)** Representative MADM clones induced at E9.5, showing multiple distinct morphotypes. **c**) Thoracic level; 400 μm projection. **d**) Cervical level; 300 μm projection. tdT and GFP hemilineages are labeled. Scale bars, 200 μm (image), 100 μm (traces). **e)** Morphotype composition of individual clones. Clones were induced at E9.5 (*n* = 21) and E10.5 (*n* = 2). Clones in **c** and **d** are indicated. Colour-code corresponds to morphotypes in **b**. **f)** Distribution of clonal heterogeneity index, *H_i_*, across all clones. **g)** Observed clonal *H_i_* compared to a null distribution using a permutation test. No significant deviation was detected (two-tailed t-test). Each dot represents a clone linked to the mean of its corresponding null distribution. Thick line denotes the group means. All clones were analyzed at E18.5. *n* indicates the number of clones. **Abbreviations:** TM, tamoxifen; E, embryonic day; GFP, green fluorescent protein; tdT, tdTomato; unidir, unidirectional; bidirec, bidirectional; triang, triangular; stell, stellate; Het, heterogeneity.

Clonally related neurons displayed divergent morphologies (**Fig. 3c-d**). Nearly all clones contained neurons with multiple morphotypes (**Fig. 3e**). To quantify morphological diversity within clones while accounting for clone size, we defined a heterogeneity index (*H_i_*) that relates the number of distinct morphotypes to the number of reconstructed neurons per clone (**Fig. 3f**). The majority of clones displayed intermediate to high morphological heterogeneity (**Fig. S7a-c**), with only one small clone containing paired sister neurons with matching morphologies (**Fig. S7a-c**).

To statistically determine whether this high degree of morphological diversity reflects a stochastic distribution of morphotypes, we performed a permutation test. For each clone, we compared the observed *H_i_*to a null distribution generated by repeatedly sampling the same number of neurons at random from the pooled population (**Fig. S7d-e**). In no case did clonal heterogeneity deviate significantly from the null expectation (**Fig. 3g**). Thus, although clonally related neurons were developmentally linked, their morphological identities appeared unconstrained by lineage.

### Individual NPs can generate excitatory and inhibitory neurons of different classes

To probe the multipotency of spinal progenitors at a molecular level, we examined the identity of clonally related neurons. We first performed iterative antibody labeling on a subset of clones induced at E10.5. Our first observation was that clones in the superficial dorsal horn contained cells expressing inhibitory dI4 markers (Pax2/Lhx5) and excitatory dI5 markers (Lmx1b and Tlx3), indicating mixed dI4/5 identity of clonally-related cells (**Fig. 4a**). Similar excitatory/inhibitory mixing was also found in a ventral clone, which was less abundant at this stage and contained clonally-related neurons expressing both inhibitory IN markers Pax2/Lhx5 and the excitatory V0 marker Evx1 (**Fig. 4b**). Furthermore, some clones contained both interneurons and motor neurons (**Fig. 4c**). These examples demonstrated that individual NPs can generate clonally-related neurons with different neurotransmitter profiles and cardinal class identity.

**Figure 4.**
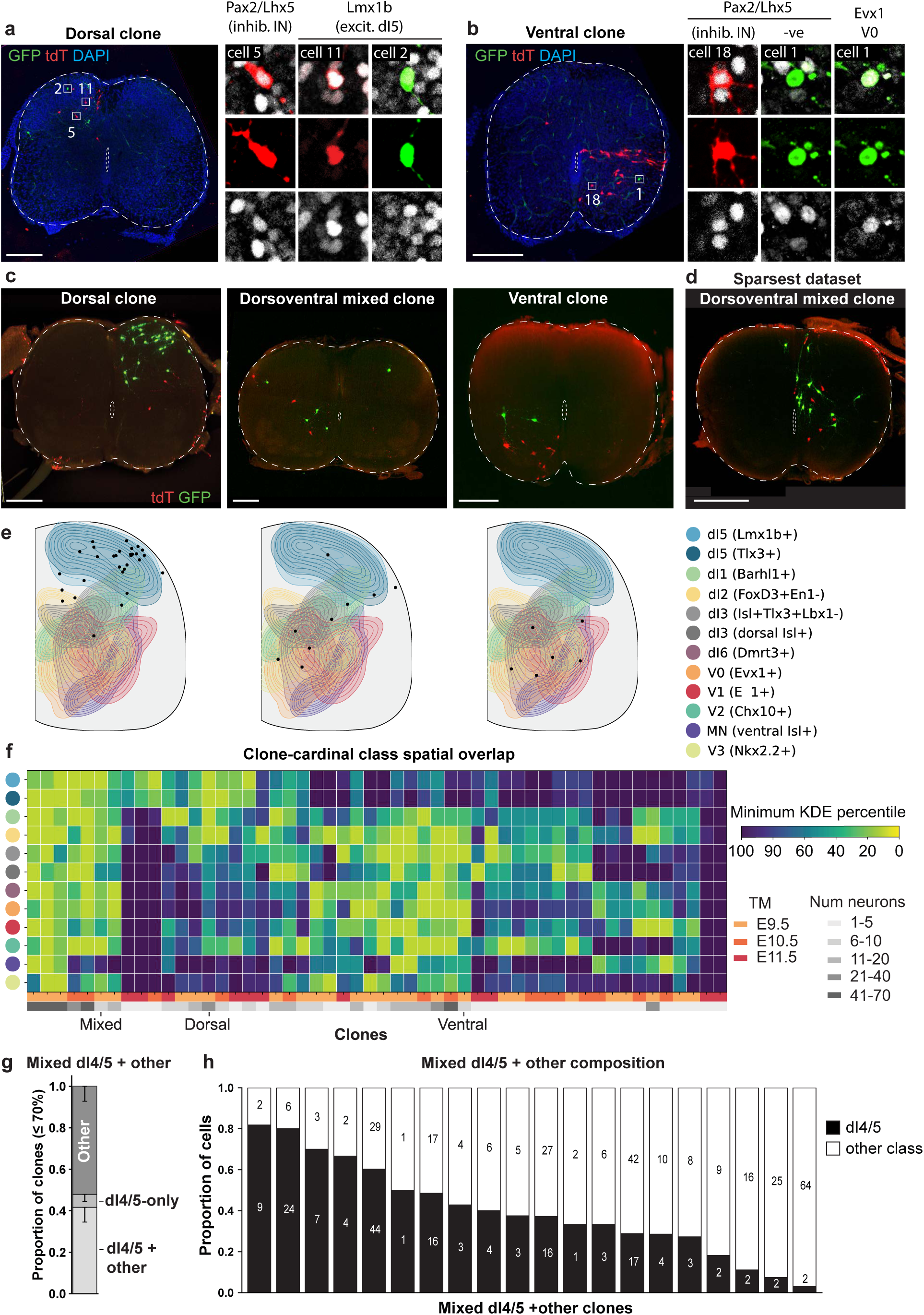
Clones contain mixed excitatory and inhibitory cardinal classes. **a)** Spinning disk confocal image of a dorsal MADM clone induced at E10.5, expressing Pax2/Lhx5 or Lmx1b. **b)** Spinning disk confocal image of a ventral MADM clone induced at E10.5, expressing Pax2/Lhx5 or Evx1. **c)** Example maximum intensity projections of clones containing dorsal (TM E9.5, lumbar level; 150 μm projection), dorsoventral mixed (TM E9.5, cervical level; 170 μm projection), or ventral neurons (TM E9.5, thoracic level; 100 μm projection). tdT and GFP hemilineages are labeled. The ventral clone contains interneurons and a motor neuron. **d)** Example of a dorsoventral mixed clone in a spinal cord very sparsely labeled with a total of four MADM events. Thoracic level; 150 μm projection. **e)** 2D Neurons positions of clones in (**c**) projected onto the kernel density estimate (KDE) maps of cardinal class marker expression distributions (see **Fig. S8**). **f)** Per-clone heatmap indicating spatial overlap of neurons with cardinal class marker distributions. Rows represent individual clones; columns correspond to cardinal classes. Heatmap values denote the lowest KDE percentile occupied by any neuron in the clone within each class distribution. Warmer colours indicate overlap with the core of the class distribution. The colour bars indicate TM injection time point and the number of neurons per clone. E9.5 (*n* = 27 clones), E10.5 (*n* = 14), E11.5 (*n* = 11). **g)** Proportion of clones containing dI4/5s and other classes, dI4/5s-only, or other classes-only based on exclusive neuron overlap with the 70th KDE percentile of class marker distributions. Proportions are presented with SE of the proportion (lower limit). Clones not fitting these categories were excluded. *n* = 48. Scale bar, 200 μm. **h)** Proportions of dI4/5s and other classes across mixed dI4/5 + other clones. All clones were analyzed at E18.5. *n* indicates the number of clones. **Abbreviations:** TM, tamoxifen; SE, standard error; E, embryonic day; GFP, green fluorescent protein; tdT, tdTomato; KDE, kernel density estimate; MADM, mosaic analysis with double markers.

Next, following the observation that many clones contained neurons occupying remote regions spanning the dorsoventral axis (**Fig. 4c-d; S9a**), we investigated whether the position of clonally related neurons could inform us about their identity. We created transverse maps of the 2D kernel density estimates (KDEs) of cardinal class spatial distributions based on immunofluorescence labeling of key transcription factors (**Fig. S8a**). These match cardinal class distributions previously characterized by genetic tracing^14^

We then projected neuron-containing clones onto these KDE maps and scored the spatial overlap of each cell with each cardinal class region, assigning predicted cardinal class membership to every neuron within the clone (**Fig. 4e; S9c**). Potential class membership based on spatial location is congruent with molecular identification in the examples above (**Fig. S9d-e**). Spatial overlap between clones and cardinal classes was then visualized in a per-clone heatmap, which depicts for each cardinal class, the highest density region (lowest KDE percentile) occupied by neurons within a clone (**Fig. 4f**). Large clones labeled at E9.5 and E10.5 frequently contained cells overlapping multiple cardinal class distributions (**Fig. 4f**). Smaller clones and those labeled at E11.5 overlapped with fewer classes (**Fig. 4f**). Because cardinal class regions overlapped with each other substantially (**Fig. S9b**), spatial information alone offered a prediction but often did not support unique class assignment.

Neurons in the superficial dorsal horn, dI5 INs (and potentially the similarly distributed dI4 population), however showed minimal spatial overlap with other classes. Therefore, we focused on distinguishing dI4/5 INs from more ventral classes. Strikingly, mixed clones of dI4/5 and ventral predicted identity remained very common (**Fig. 4f**); nearly ∼45% of clones met the criterion for containing superficial dorsal horn (dI4/5-like) cells together with at least one ventral cardinal class (**Fig. 4g-h**). These findings revealed that individual NPs can generate neurons spanning broad dorsoventral territories, including both superficial and deep spinal laminae, and include multiple cell types.

## DISCUSSION

Spatiotemporal patterning in the spinal cord instructs neuron identity through progenitor location and time of birth^10^. Here, single-progenitor lineage tracing in the spinal cord extends this canonical view by revealing that spinal progenitors are profoundly multipotent, defying a simple model of cardinal class-dedicated progenitors. Single lineages produce both neurons and glia in highly variable numbers, contain neurons of diverse cell-type identity and morphology, and span sensory-motor territories and even the two hemispheres of the spinal cord. Surprisingly, lineage heterogeneity thus emerges as a defining principle of spinal cord development.

### Bilateral lineage dispersion and midline crossing

While spinal cord development is classically framed in terms of neurogenesis and fate specification on the same side, or in a hemicord, our lineage tracing reveals an, unanticipated dimension of its development: bilateral dispersion within a clone. Strikingly, ∼40% of clones contribute cells to both hemispheres of the spinal cord. Mechanistically, this raises the question of whether progenitors or post-mitotic cells cross the midline. Several lines of evidence argue against progenitor dispersal as the mechanism of this bilaterality. Previous studies show dispersal is limited in the neural tube^23,39^ and lumen forms a physical barrier along most of the DV midline until partially fusing at E15^40,41^. Furthermore, many clones display bilaterality in both MADM color-coded hemilineages, which would require not one but two separate progenitor crossing events.

Migration of post-mitotic cells across the CNS midline is unusual but has been observed by neurons in the mouse hindbrain^42–48^ and by glia in rare events in the chick spinal cord^18,19^. In the hindbrain, commissural oculomotor and precerebellar neuron populations form their contralateral projections via soma migration across the ventral midline. Similarly, bilateral clones in the spinal cord are enriched in commissural neurons, including predicted dI1, dI2, dI4s, V2a, and V3 interneuron classes^49,50^. Well-established mechanisms of axon guidance have been defined for many of these spinal commissural populations^51^. However, they do not rule out the possibility of soma migration. Consistent with such a soma migration hypothesis, within a bilateral spinal cord clone, contralateral neurons are often found at the same dorsoventral level as the ventral or dorsal commissure, even seemingly connected by a commissural axon tract in one example. Such observations raise further questions about the existence, mechanism, and purpose of soma migration in the spinal cord, including whether somas can translocate along existing axon tracts, whether reciprocal contralateral connections are formed between sister neurons, and whether these commissural populations comprise a specialized, functional cell type.

### Highly variable, non-stereotyped division patterns for spinal lineages

Cell division in growing organs must be regulated to ensure appropriate size and composition. In the mouse cerebral cortex, radial glial cells progress through a stereotyped lineage progression, undergoing predictable numbers of proliferative and neurogenic divisions and producing units of 8-9 neurons^26^. In contrast, spinal progenitors undergo variable rounds of proliferation, do not produce a unitary output of neurons or glia, and lack an apparent concerted temporal lineage progression. Similar to systems like the retina and superior colliculus^4,52^, cell proportions in the spinal cord may be balanced at the population level rather than regulated through stereotyped lineages. In the spinal cord, such variability may accommodate rostrocaudal level-specific differences in neuronal composition or balance cell-type proportions over the extended window of neurogenesis.

The lineage variability that we observe does not preclude however that domain-specific patterns in progenitor behavior also occur. Such patterns are supported by live imaging of cell division in the zebrafish spinal cord, in which distinct modes of neurogenesis generate ventral IN subtypes in the p0 and p2 domains^53,54^, with V2a/b interneurons generated via terminal, Notch-differentiated divisions. Such stereotyped p2 neurogenic behavior could occur within a larger clone or be difficult to resolve in our dataset because clones span all dorsoventral domains. Future analysis has the potential to reveal such finer-scale, progenitor-specific patterns.

### Lineage divergence in the context of domain patterning

In many tissues, different types of neurons are generated in a stereotyped sequence by multipotent progenitors with temporal changes in competence^55,56^. Consistent with this, our data demonstrate that the dorsoventral spread of a clone in the spinal cord decreases over time, making later-born, clonally-related neurons are more likely to be of a similar type. However, the dramatic mixing of putative dI4/5 with more ventrally settling interneuron classes that we observe in earlier-born clones strongly supports high levels of progenitor multipotency beyond a simple one-progenitor-to-one-cardinal-class model.

Other lineage tracing work in the spinal cord has also suggested mixing of cardinal classes. Pioneering retroviral tracing in the chick described motor neuron-containing clones which also contained ventral or even dorsal interneurons^18^. Recent work in the Briscoe lab finds mixing of spinal cord neuron types within five lineage subdivisions, each containing multiple cardinal classes generated by neighboring progenitor domains^58^. Some of the dorsal-ventral mixing we observe within a clone could be explained via similar identity switches across domain boundaries. For instance, the juxtaposed dorsal progenitor domains dp5 and dp6 generate the dorsal-most dI5 and ventral dI6 interneurons, respectively. However, such mixing cannot account for motor and inter-neuron containing clones, which we and others^18^ observed.

Such extreme diversity and dispersion of clonally related neurons implies that early progenitor patterning is not fully sufficient to predict the ultimate identity of post-mitotic neurons. Additional fate allocation mechanisms may contribute to the variability in clone composition^59^. In the CNS, such mechanisms can include the apparent stochasticity in neurogenic decisions that contribute to diversifying cortical and retinal clones^4,7^ and/or target-derived secreted signals or electrical activity that have been shown to regulate neuron fate in the spinal cord and cortex, including the neurotransmitter expression, migration, and motor pool-specific position and connectivity of spinal motor neurons in mice and frogs^61–65^. Alignment of the position of clonally-related neurons with either anatomical-projection maps or spatial transcriptomic atlases may give further insight into target-derived cues or shared gene expression programs driving more dramatic intraclonal neuron dispersion.

### Shared neuronal and glial lineages

The presence of multipotent neuronal-glial progenitors in the spinal cord has long been debated^13^. Our findings reveal that most spinal progenitors undergo gliogenesis, generating several types of astrocytes and other glia. Contrasting previous results^66,67^ we do not find evidence of early specification of a glial-dedicated progenitor in the spinal cord: neurons always co-occured with glia before E11.5. However, as not all cell types were identified within every clone based on morphology, we cannot exclude that there exist rare or later-born glia-only clones. Our data also aligns with the gliogenic output that was previously observed for the p0-p3 domains, for which TFs involved in neurogenic fate specification programs are repurposed to generate astrocytes in chick and mice^68–70^. Furthermore, we find that this high prevalence of mixed neuron-astrocyte clones spans the entire dorsoventral axis, supporting that astrocytes are generated by not one but multiple, and perhaps all, progenitor domains. Mixed output is thus rather the rule than the exception. Such diversity in glial origin suggests diversity in function, as astrocytes derived from different regions of the mouse spinal ventricular zone are positionally distinct^71,72^ and astrocytes with different ontogenies have different reactivity^73^. Lineage origin thus emerges as a potential driver of functional specialization for spinal cord glia^13^.

### Lineage versus function

Beyond influencing cell fate, intrinsic lineage programs can also contribute to the functional organization of neural tissues. In the *Drosophila* nervous system and the mammalian cortex and retina, progenitors produce sets of neurons that together comprise a functional unit^1^. In the vertebrate brain, lineage influences dendritic morphology and connectivity, with sister neurons often participating in shared circuits^74–81^. By contrast, V2a/b sister interneurons in zebrafish share a gross morphology but integrate into separate microcircuits^82^.

We observed little obvious correspondence between the clonal architecture and functional organization of the spinal cord, based on spatial, morphological, or molecular properties. Clonally related spinal neurons frequently dispersed across dorsal sensory and ventral motor regions and were not confined to specific Rexed laminae. Unbiased analyses of proximal spinal neuron morphology showed lineage relationships do not constrain neuronal morphology. Moreover, individual progenitors generate both excitatory and inhibitory interneurons, and even mixed motor/interneuron and interneuron cardinal identities. Together, the broad settling positions, morphological heterogeneity, and mixed subtype, cardinal class and neurotransmitter profiles of clonally related neurons in the mouse spinal cord suggest that, unlike in the brain, clonally related spinal neurons may participate in distinct spinal circuits, or circuits that are linked by a functional axis that is still unknown.

Together, our results thus point to a new, previously unappreciated aspect of spinal cord development in which progenitor domains bias, but do not rigidly determine, neuronal fate. Instead, individual and highly multipotent progenitors generate variable numbers of neurons and glia that disperse widely, and even bilaterally, and are poised to integrate into functionally distinct circuits.

## RESOURCE AVAILABILITY

### Lead contact

Further information and requests for resources and reagents should be directed to and will be fulfilled by the lead contact, Lora B. Sweeney.

### Materials availability

All materials will be made available upon request to the lead contact.

### Data and code availability

The code used for clonal and morphological analyses is available at Github: https://github.com/sgobeil1/SCLineage

## ACKNOWLEDGMENTS

We would like to thank Elizabeth Marin, Anna Kicheva, Igor Adameyko, and James Briscoe as well as members of the Sweeney and Hippemeyer labs and SFB consortium for comments on the manuscript. We are also grateful for the technical support of the Preclinical and Imaging and Optics Facilities support teams (ISTA). In addition, we thank our funding sources for providing the resources to do these experiments: Horizon Europe ERC Starting Grant Number 101041551 (M.S.; L.B.S.); Special Research Program (SFB) of the Austrian Science Fund (FWF) NeuroStem Modulation Project numbers F7814-B (S.A.G.; M.S.; G.S.; and L.B.S.) and F7805 (G.C. and S.H.). S.A.G is supported by a Boehringer Ingelheim Fonds PhD Fellowship, F.D.S.N. by an Institute of Science and Technology Austria (ISTA) GROW fellowship, and G.C. by an ISTA Plus postdoctoral fellowship from the European Commission. S.H./L.B.S. and G.C. were additionally supported by institutional funds from the ISTA and the University of Exeter, respectively.

## AUTHOR CONTRIBUTIONS

L.B.S. led and coordinated the project. S.A.G., S.H., and L.B.S. conceived of the study and designed experiments. S.A.G., with support and training from G.C. and C.S., performed all experiments. S.A.G., F.S.N., G.S., and M.S. quantified the data. S.A.G. and F.S.N/S.A.G. designed and implemented analysis pipelines and software for clonal analysis and morphological analysis, respectively. S.A.G., F.S.N., and L.B.S. wrote the manuscript; S.H. and G.C. edited the manuscript and provided critical feedback.

## DECLARATION OF GENERATIVE AI AND AI-ASSISTED TECHNOLOGIES

During the preparation of this work, the authors used ChatGPT 5.2 and DeepsSeek-V3.2 to brainstorm alternative, more concise language, and coding solutions. After using these tools, the authors reviewed and edited the content as needed and take full responsibility for the content of the publication.

## SUPPLEMENTAL INFORMATION

Extended Data including Supplementary Figures 1-9.

## Methods

### Mice

All animal procedures were approved by the Austrian Federal Ministry of Science and Research and carried out in compliance with the Austrian and European Union animal law (license numbers: 2024-0.191.328 and 2025-0.146.987). Mice were bred and maintained under protocols approved by the relevant institutional animal-care and ethics committees and in accordance with guidelines of the preclinical facility (PCF) at the Institute of Science and Technology Austria. Animals were housed under controlled conditions (21LJ±LJ1LJ°C; 40–55% humidity) on a 12 h light/12 h dark cycle. The transgenic lines carrying MADM cassettes on chromosome 11^84^ and *S*ox*2^CreER^* ^83^ have been described previously. All strains were maintained on a mixed C57BL/6, FVB, and CD1 background; wild-type CD1 mice were additionally used in select experiments. Both sexes were included, using littermates of the appropriate genotypes. Animal numbers were minimized wherever possible in line with the 3Rs (replacement, reduction, refinement).

### Generation of Experimental Mice

MADM clones were induced in the spinal cord as previously described^26,27,39^. *MADM-11^GT/TG^;S*ox*2^CreER^* mice were generated by crossing *MADM-11^GT/GT^;Sox2^CreER^* with *MADM-11^TG/TG^* mice. To induce MADM clones, timed pregnant females were injected intraperitoneally with tamoxifen (TM; 0.5-2 mg/mouse dissolved in corn oil) at E9.5, 10.5, or 11.5. At E18.5, litters were delivered by caesarean section.

### Tissue collection and processing

E18.5 embryos were anesthetized by inhalation of isoflurane and transcardially perfused with 3-5 mL of ice-cold 4% paraformaldehyde (PFA), then fixed overnight in 4% PFA. Spinal cords were dissected and cut into six pieces corresponding to rostrocaudal level based on gross morphology of the cervical and lumbar enlargements: two cervical, two thoracic, one lumbar, and one lumbosacral. The spinal cords were then cleared for wholemount or cryosectioned tissue processing (see below).

### Wholemount tissue processing

#### Tissue clearing

Spinal cords were embedded in 1% agarose (dissolved in H_2_O) before clearing. The clearing protocol was adapted from published CUBIC-clearing protocols^85,86^. Embedded spinal cords were washed in 50% 1:1 CUBIC-L:CUBIC-R1a solution in H_2_O (CUBIC-L: 10% w/v Triton X-100, 10% w/v N-butyldiethanolamine in dH_2_O; CUBIC-R1a: 10% w/v urea, 5% w/v N,N,N’,N’-tetrakis, 10% w/v Triton X-100, 1:200 5M NaCl in dH_2_O) for 6h at 37°C, 300 rpm. Then, spinal cords were incubated in 100% 1:1 CUBIC-L:CUBIC-R1a solution for 48h at 37°C, 300 rpm, with one solution change. Spinal cords were then washed twice with PBS for 2 hours then once overnight at 37°C on a rotator at 300 rpm. Immunolabeling (see below) was then performed at this point. After washing off the secondary antibody, the refractive index of the cleared spinal cord was matched by stepwise transfer into CUBIC-R+(N) (45% Antipyrine, 30% Nicotinamide in dH_2_O): two-hour steps of 25%, 50%, 75%, 100% CUBIC-R+(N) in dH_2_O were performed. The CUBIC-R+(N) was replaced twice, including at least one overnight incubation, before imaging.

#### Immunolabeling

Cleared spinal cords were incubated in GFP and tdT primary antibodies (see **Table 1**) diluted in 0.2% PBST (0.2% Triton X-100 in PBS) for 4 days at 37°C, 300 rpm. The antibody solution was removed and cords were washed three times for 2 hours with PBS at 37°C on a rotator at 300 rpm. Spinal cords were next incubated with secondary antibodies (see **Table 2**) diluted in 0.2% PBST for 2 hours at 37°C again on a rotator at 300 rpm. The antibody solution was removed and cords washed once for 2 hours with PBS at 37°C, again rotating at 300 rpm, before proceeding with refractive index matching.

**Table 1.** Primary antibodies list.

| Target | Dilution | Species | ID | Lot or bleed date | Source |
| --- | --- | --- | --- | --- | --- |
| Barhl1 | 1:1k | Rabbit | HPA004809 | 000037963 | Sigma Aldrich |
| Chx10/Vsx2 | 1:500 | Sheep | AB16141 | 1013906-1,<br>1070814-4 | Abcam |
| Dmrt3 | 1:32k | Rabbit | CU1078 | NA | Columbia University |
| DsRed | 1:16k | Guinea pig | CU1906 | 7/14/2017 | Columbia University |
| En1 | 1:16k | Guinea pig | CU1714 | 6/08/2012 | Columbia University |
| Evx1 | 1:400 | Mouse (IgG2a) | 99.1.3A2 (supernatant) | 5/12/2022 | DSHB |
| Foxd3 | 1:32k | Rabbit | NA | Terminal bleed | Carmen Birchmeier |
| GFP | 1:2k | Chicken | ab13970 | 1018753-23,<br>1018753-31 | Abcam |
| Isl1/2 | 1:25 | Mouse (IgG2b) | 4D5-16-1 | 08/04/2023 | Columbia University |
| Lbx1 | 1:32k | Guinea Pig | NA | Terminal bleed | Carmen Birchmeier |
| Lhx5 | 1:2k | Rabbit | T3a | NA | Columbia University |
| Lmx1b | 1:32k | Guinea Pig | NA | Terminal bleed | Carmen Birchmeier |
| Nkx2.2 | 1:500 | Mouse (IgG2b) | 74.5A5 (supernatant) | 8/4/2022 | DSHB |
| Pax2 | 1:1k | Rabbit | 71-6000 | WA317232 | Invitrogen |
| RFP | 1:1k | Rabbit | PM005 | 049 | MBL |
| Sox9 | 1:100 | Rabbit | 82630 (D8G8H) | NA | Cell Signalling Technology |
| tdTomato | 1:500 | Goat | AB8181 | 81260924 | Sicgen Antibodies |
| Tlx3 | 1:32k | Rabbit | NA | Terminal bleed | Carmen Birchmeier |

**Table 2.** Secondary antibodies list.

| Target | Fluorophore | ID | Lot | Source |
| --- | --- | --- | --- | --- |
| Chicken | Alexa fluor 488 | 703-545-155 | 165794, 170532, 175023, 168728 | Jackson Immunoresearch |
| Guinea pig | Cy5 | 706-175-148 | 175232, 170461 | Jackson Immunoresearch |
| Guinea pig | Cy3 | 706-165-148 | 173590, 172797 | Jackson Immunoresearch |
| Guinea pig | Alexa fluor 405 | ab175678 | 1107206-5 | Abcam |
| Mouse - IgG2a | Alexa fluor 488 | 115-545-206 | 167697 | Jackson Immunoresearch |
| Mouse - IgG2a | DyLight 405 | 115-475-206 | 167175 | Jackson Immunoresearch |
| Mouse - IgG2b | Cy5 | 115-175-207 | 156313 | Jackson Immunoresearch |
| Mouse - IgG2b | DyLight 405 | 115-475-207 | 175714 | Jackson Immunoresearch |
| Goat | Cy3 | 705-165-147 | 174616,172428 | Jackson Immunoresearch |
| Rabbit | Cy5 | 711-175-152 | 171768 | Jackson Immunoresearch |
| Rabbit | Cy3 | 711-165-152 | 175018, 173690, 172096 | Jackson Immunoresearch |
| Rabbit | Alexa fluor 405 | ab175649 | 1092037-5 | Abcam |
| Sheep | Cy3 | 713-165-147 | 168633,168633 | Jackson Immunoresearch |

#### Image acquisition

Cleared spinal cords embedded in 1% agarose were imaged with a Zeiss Lightsheet 7 using a 5x/0.16 NA objective and 488 and 561 nm excitation lasers. CUBIC-R+(N) or mineral oil with a matching refractive index of 1.52 were used as immersion media. Zeiss image files were converted to Imaris file format and analyzed as described in Clonal Analysis and Quantification.

### Cryosectioned tissue processing

#### Cryosectioning

Spinal cords were cryoprotected by incubating in 30% sucrose solution (30% sucrose in 1:1 0.2M phosphate buffer:H_2_O) overnight at 4°C, rotating at 70 rpm. Next, spinal cords were embedded in optimal cutting temperature compound (O.C.T., Tissue-Tek), frozen, cryosectioned in the transverse plane at 50 μm, and mounted on Superfrost Plus slides.

#### Immunolabeling

Antibody labeling of key TF factors to map cardinal class distributions was performed on sectioned E18.5 CD1 or *MADM-11^GT/TG^* spinal cords. Slides were washed once with PBS then incubated with primary antibodies (see Table 1) diluted in 0.2% PBST for 2 days at 4°C. The antibody solution was removed and slides were washed three times with PBS. Next, slides were incubated in secondary antibodies (see Table 2) diluted in 0.2% PBST for 4 hours at room temperature. The antibody solution was again removed and slides were washed three times with PBS for 10 min, and then coverslipped with homemade Mowiol-DABCO mounting medium (25% w/v glycerol, 10% w/v Mowiol, 2.5% w/v DABCO in 0.2 M Tris-HCl pH 8.5).

#### Iterative immunolabeling

Molecular expression analysis of MADM clones was performed using an iterative antibody labeling protocol adapted from published protocols^87,88^. Sectioned spinal cords were post-fixed with 4% PFA at room temperature for 15 minutes to improve section adherence. Immunolabeling was carried out as described above. Briefly, after washing off the secondary solution, slides were coverslipped with imaging buffer (700 mM N-acetyl cysteine, 10% glycerol in H_2_O, pH adjusted to 7.4). Slides were imaged as described below. After imaging, slides were gently decoverslipped by placing them in a vertical staining jar filled with PBS and allowing the coverslips to slide off. Slides were washed four times with PBS. Antibodies were eluted by incubation in elution buffer (0.64 M L-glycine, 3 M urea, 3 M guanidinum chloride, 70 mM TCEP in H_2_O, pH adjusted to 2.5) twice for 5 min at room temperature. After three PBS washes, slides were stored in PBS at 4°C or incubated in primary antibody solution to begin the next round of labeling.

#### Image acquisition and processing

Immunolabeled clones were imaged at a spinning disk confocal microscope (Yokogawa CSU-W1 dual-disk spinning disk, 50 μm pinhole, with Nikon Ti2E) using a 20x air/0.75 NA objective and 405nm, 488nm, 561nm, 640nm excitation lasers. Immunolabeled sections for the cardinal class distribution map were imaged at the spinning disk or confocal microscope (Zeiss Axio Observer Z1 Inverted LSM900, and Axio Imager.Z2 Inverted LSM800 confocal microscopes) using a 20x air/0.8 NA objective and 405 nm, 488nm, 561nm, and 640nm excitation lasers. Nikon and Zeiss files were converted to Imaris file format and analyzed as described in Cardinal class distributions.

### Antibodies

#### Clonal Analysis and Quantification

Cell coordinate tables were exported from Imaris and analyzed using a custom Python pipeline. Briefly, cell coordinates were normalized to the central canal (0,0) and to an average E18.5 spinal cord rostrocaudal length (1.47 cm). Clones of red/green cells were defined based on inter-cell and inter-clone thresholds (see below **Clonal resolution**). Quality-control exclusion criteria were applied to remove clones with tissue damage (caused during dissection), close proximity to sample preparation cut sites (within 100 μm), and ambiguous proximity to other cells (see below **Clonal resolution**). Unless otherwise stated, single-cell clones were excluded from clone quantifications. Clones containing cells of only one hemilineage, indicating apoptosis of the other hemilineage, were excluded from quantifications of progenitor cell division patterns.

#### Clonal resolution

Sparse MADM labeling is essential to achieve clonal resolution. Consistent with previous MADM studies^26,39,52^, several lines of evidence supported the clonality of our approach. First, discrete clusters of red/green (R/G) or yellow cells were distributed along the rostrocaudal (RC) axis of the spinal cords (**Fig. 1c,d; S1a,c**). Second, the RC nearest-neighbor distance (NND) among R/G and among yellow cells were small, reflecting their tendency to cluster among themselves (**Fig. S1d**). Third, and in contrast, the NND between R/G and yellow cells was larger than in a simulated random data set. These results support that the discrete clusters of labeled cells we observed are clones originating from separate progenitors.

To further confirm clonal resolution, we assessed clonal architecture in samples with particularly sparse labeling (≤5 labeling events per spinal cord) and further narrowed our criteria for stringent clone identification in the complete sample set (**Fig. S1e-j**). Cells located within 125 μm along the rostrocaudal axis were considered part of the same clone, while distinct clones were required to be separated by at least 200 μm. On average, identified clones were spaced tenfold farther than this minimum threshold (**Fig. S1i–j**). Cell clusters that did not meet these criteria were excluded to eliminate ambiguity regarding clonality.

#### Nearest-neighbor distance (NND) analysis

Nearest-neighbor distance (NND) analysis was performed as previously described^26^ by computing, for each cell, the Euclidean distance to its closest neighboring cell in 3D space (XYZ). An expected distribution under spatial randomness was generated by simulating the same number of points within a volume matched to the average sample dimensions 100 times, repeating the NND analysis on the simulated points.

#### Cell-type identifications based on morphology

The following cell types were identified based on their morphology: *neurons*, cells with recognizable neurites; *motor neurons*, neurons located in the large soma regions of the MMC or LMC and possessing extensive arborizations; *commissural neurons*, neurons possessing a neurite that crosses the midline; *septal cells*, cells positioned on the dorsal or ventral midline with vertical processes spanning the ventricular to pial surface; *protoplasmic astrocytes*, grey or white matter cells with star-like dense network of highly branched, often indistinct processes; *fibrous astrocytes*, white matter cells with elongated, often dense, processes; *glia with endfeet*, astrocyte-like glial cells which extend a radial process to an endfoot on the pial surface; *glial limitans astrocytes*, cells with somas just under the pia (these are distinguished from glial endfeet by their size and absence of attachment to a soma in the parenchyma); *progenitors*, cells adjacent to the central canal without neurites; *other glia*, white matter cell with ambiguous morphology. Cells with *uncertain* identity that could not be classified into one of these groups due to ambiguous morphology or lack of imaging clarity, were excluded from cell-type quantifications (26.8% of all red/green cells).

### Spatial analysis of MADM clones

#### Cardinal class distributions

Cardinal class transverse distributions, defined by transcription factor (TF) marker expression, were generated using kernel density estimations (KDEs). Coordinates of cell nuclei labeled with the respective marker antibodies were recorded using Imaris. For marker co-expression counts, a custom MATLAB script was used to identify colocalized nuclei within a 4.5 μm radius. The XY coordinates of the cells were normalized to a reference lumbar hemisection using four anatomical reference points (central canal, most ventral point of the white matter, most lateral point of the white matter, and most dorsal point of the white matter) using custom MATLAB and Python scripts modified from previous work^89^. These marker-expressing cel^89^populations were converted into 2D kernel density estimates (KDEs) on a common fixed rectilinear grid. KDEs were generated using Gaussian KDE bandwidth selection (Scott’s rule) and normalized to integrate to ∼1 over the grid. For each reference KDE, we computed highest-density region (HDR) thresholds at the 10th-70th percentiles. The intersection areas of the 70th percentile HDRs of pairs of KDEs were calculated using the Jaccard index (overlap area/union area).

#### Predicting cardinal class identity of neurons within a clone

Clones with a minimum of 80% of morphologically identified cell types were included in the analysis. The XY coordinates of neurons within these clones were normalized to the reference hemisection using the four anatomical reference points in a similar way. Coordinates from both hemispheres were mirrored into one. Each pair of coordinates was then evaluated against each cardinal class reference KDE using bilinear interpolation on the fixed grid and assigned the *smallest* HDR percentile it satisfied (e.g., “≥10%”, “≥20%”, …), or “Outside” if it fell outside all queried HDRs. The resulting per-neuron KDE/HDR membership table was used to generate heatmaps showing membership per clone or per cell. To quantify “mixing” of dI4/5s and other cardinal classes, clones were scored by exclusive membership of at least one neuron in the 70th percentile HDR of dI5s and at least one neuron in any other cardinal class.

### Morphological analysis

#### Reconstruction of neuronal morphology

Clones with optimal clearing and image quality were included in the morphological analysis. Neurons were included in the analysis if their total reconstructed neurite length exceeded 400 µm. A minimum of two neurons were reconstructed per clone. Overall, an average of 55% of neurons per clone were reconstructed. MADM-labeled neurites were manually traced in 3D using Imaris (9.9.1). Tracing was initiated at the soma surface and continued until the neurite terminated, became too faint to reliably trace, or entered a region of dense labeled cells where individual neurites could not be distinguished. Reconstructed neurons were exported in SWC file format.

#### Quantitative analysis of proximal neuronal morphology

Proximal neuronal morphology, within a 100 μm radius of the soma, was analyzed. Similar to previous work, arborization complexity peaked within the 100 μm radius^37^. Three morphological feature^37^were quantified: (1) number of primary neurites, (2) neurite geometry, and (3) branching complexity.

Primary neurites were defined as neurites emerging directly from the soma. This information is recorded in the SWC files.

Neurite geometry was assessed by measuring the angles between primary neurites. For each neurite, a vector was measured from the soma to the neurite intersection with a 50 μm radius or, if they were reached sooner, the first branching point or the neurite termination point. The neurite vectors were projected onto the transverse plane and the angular gaps between neighboring neurites were calculated. The angles and ratio of the largest and second-largest gaps, in addition to the number of primary neurites, were used to classify neurons based on the geometrical configuration of their primary neurites^37^ (**Fig.** 3**b**).

Branching complexity was quantified using Sholl analysis as the total number of neurite intercepts at concentric 2 μm intervals within the 100 μm radius. This metric is consistent with approaches used for proximal morphological assessment in spinal interneurons^38^. For each geometric category, a branching complexity threshold was set at the mean total intercept number. Neurons with total intercepts above this threshold were categorized as “complex” and those below “simple”. This relative classification accounts for inherent differences in branching potential across configurations. Combining five geometries with two complexity levels yielded ten morphotypes that covered all reconstructed neurons.

#### Statistical analysis of clone morphological heterogeneity

For each clone, a heterogeneity index, *H_i_*, was calculated to quantify the number of morphotypes and account for variability in the number of reconstructed neurons:

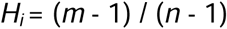

Where *m* is the number of morphotypes and *n* is the number of reconstructed neurons in the clone. This index varies from 0 (all neurons classified as one morphotype) to 1 (every neuron classified as a distinct morphotype).

To determine whether observed levels of clonal morphological diversity were greater than would be expected by chance, a permutation test was conducted. For each clone size *n* (number of reconstructed neurons), 10,000 pseudoclones were randomly generated by drawing *n* morphotypes from a pool of all the reconstructed neurons across clones (with replacement). In this way, a null distribution of the *H_i_* for each clone, under the null hypothesis that lineage does not impose any morphological restriction, was obtained. The observed clone *H_i_*s were then compared to their null distributions to obtain a *p* value. None of the observed values significantly deviated from their null distributions.

The observed morphotypes per clone were compared to extreme models of lineage restriction and diversity by generating hypothetical clones of the same size with complete homogeneity (*H_i_* = 0), complete heterogeneity (*H_i_* = 1), or a random composition (mean of the null distribution).

## Statistical information

Clonal analysis data and immunolabeling quantification were processed using custom Python scripts and exported as comma-separated (.csv) spreadsheets for archiving and downstream analysis. Statistical tests were carried out in GraphPad Prism (v10.2.2). Data are reported as means or proportions with error bars representing the standard error of the mean (SEM) or, for proportions, the lower bound of the standard error; n denotes the number of clones unless indicated otherwise. Normality was assessed using the D’Agostino–Pearson test. Comparisons between two groups were performed using two-tailed Student’s *t* tests or Mann–Whitney tests, as appropriate. For analyses involving more than two groups, we used Kruskal–Wallis tests followed by Dunn’s post hoc multiple-comparisons correction. Differences in proportions across conditions were evaluated using Fisher’s exact test. Sample sizes, statistical tests, and exact *p* values are provided in the relevant figure legends.

