## Supplemental Information for "Lineage origin of spinal cord cell type diversity"

### EXTENDED DATA

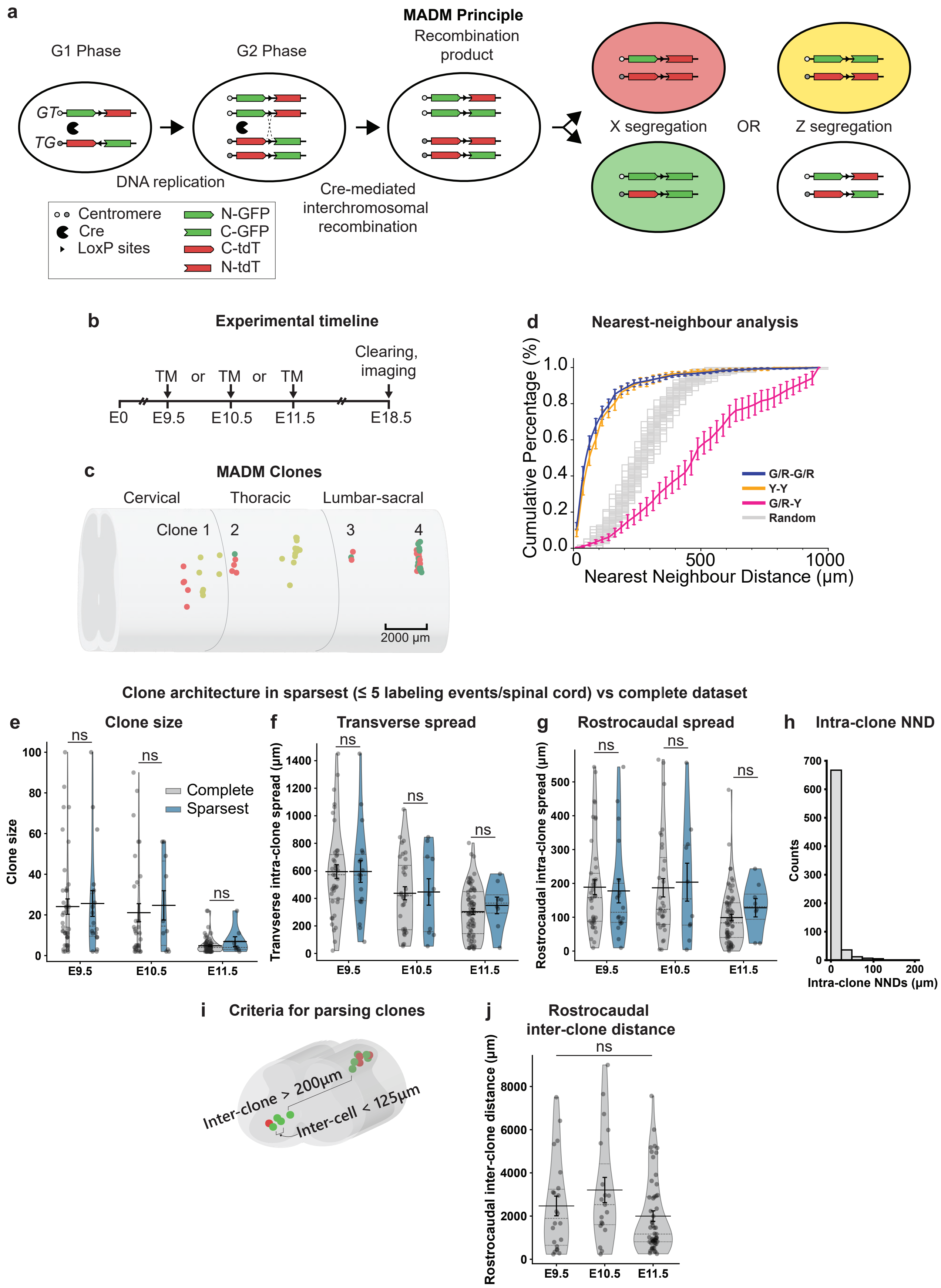
a
MADM Principle
G1 Phase
G2 Phase
Recombination product
GT
TG
DNA replication
Cre-mediated interchromosomal recombination
X segregation
OR
Z segregation
Centromere
Cre
LoxP sites
N-GFP
C-GFP
C-tdT
N-tdT
b
Experimental timeline
TM or TM or TM
Clearing, imaging
E0
E9.5
E10.5
E11.5
E18.5
c
MADM Clones
Cervical
Thoracic
Lumbar-sacral
Clone 1
2
3
4
2000 µm
d
Nearest-neighbour analysis
Cumulative Percentage (%)
Nearest Neighbour Distance (µm)
G/R-G/R
Y-Y
G/R-Y
Random
Clone architecture in sparsest (≤ 5 labeling events/spinal cord) vs complete dataset
e
Clone size
Complete
Sparsest
ns
E9.5
E10.5
E11.5
f
Transverse spread
Tranverse intra-clone spread (µm)
g
Rostrocaudal spread
Rostrocaudal intra-clone spread (µm)
h
Intra-clone NND
Counts
Intra-clone NNDs (µm)
i
Criteria for parsing clones
Inter-clone > 200µm
Inter-cell < 125µm
j
Rostrocaudal inter-clone distance
Rostrocaudal inter-clone distance (µm)

**Supplementary Figure 1. Sparse MADM clone labeling in the embryonic mouse spinal cord.**
**a)** Schematic of MADM lineage tracing.
**b)** Experimental timeline.
**c)** Example MADM-labeled spinal cord from an embryo induced at E9.5 with four clusters of red/green cells (clones) and three clusters of yellow cells. Same spinal cord as in **Fig. 1c**.
**d)** NND analysis of MADM-labeled spinal cells following induction at E9.5, E10.5, and E11.5. Cumulative density functions are presented as mean ± SEM.
**e)** Size of clones (total number of cells) across induction time points in the complete dataset vs sparsest animals ($\leqslant$5 labeling events/spinal cord). E9.5 ($n$ = 41 complete; 18 sparsest), E10.5 ($n$ = 32 complete; 10 sparsest), E11.5 ($n$ = 72 complete; 8 sparsest). Mann-Whitney test; $p$ = 0.9057, $p$ = 0.6774, $p$ = 0.4671 at E9.5, E10.5, and E11.5, respectively.
**f)** Quantification of transverse spread of clones across induction time points in the complete dataset vs sparsest animals ($\leq$ 5 labeling events/spinal cord). Mann-Whitney test; $p$ = 0.9642, $p$ = 0.8673, $p$ = 0.4321 at E9.5, E10.5, and E11.5, respectively.
**g)** Quantification of rostrocaudal spread of clones across induction time points in the complete dataset vs sparsest animals ($\leq$ 5 labeling events/spinal cord). Mann-Whitney test; $p$ = 0.6506, $p$ = 0.9826, $p$ = 0.1993 at E9.5, E10.5, and E11.5, respectively.
**h)** Frequency of NND between cells within clones in the sparsest dataset. Maximum NND is 125 µm. $n$ = 36.
**i)** Criteria for parsing clones in the complete dataset.
**j)** Rostrocaudal inter-clone distance across induction time points. E9.5 ($n$ = 41), E10.5 ($n$ = 30), E11.5 ($n$ = 72). Kruskal-Wallis test; $p$ = 0.1456.
Violin plot data are presented as mean ± SEM. All clones were analyzed at E18.5. $n$ indicates the number of clones.

**Abbreviations:** TM, tamoxifen; SEM, standard error of the mean; E, embryonic day; GFP, green fluorescent protein; tdT, tdTomato; MADM, mosaic analysis with double markers; NND, nearest-neighbor distance.

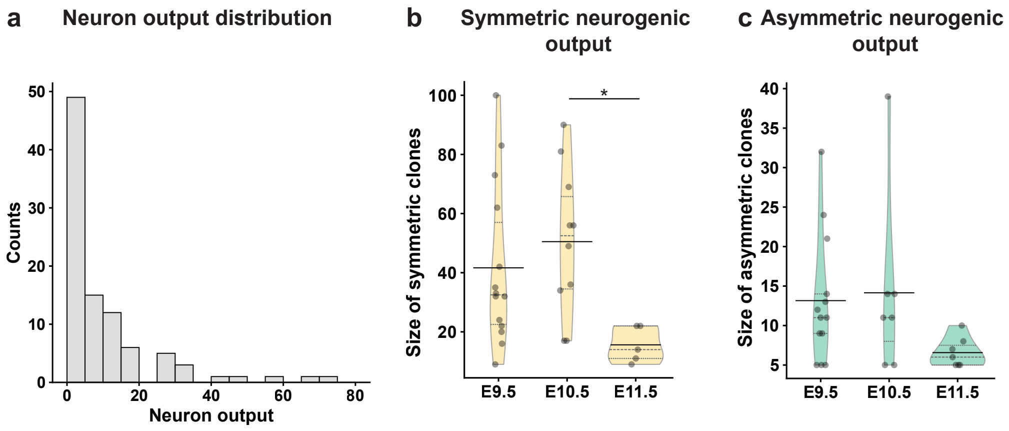

a
Neuron output distribution
Counts
Neuron output
b
Symmetric neurogenic output
Size of symmetric clones
*
E9.5
E10.5
E11.5
c
Asymmetric neurogenic output
Size of asymmetric clones
E9.5
E10.5
E11.5

**Supplementary Figure 2. Symmetric and asymmetric MADM clone output.**
**a)** Distribution of neuron output (number of neurons per clone) across all neurogenic clones. $n$ = 95.
**b)** Size of symmetric neurogenic clones (number of cells) across induction time points. E9.5 ($n$ = 14), E10.5 ($n$ = 10), E11.5 ($n$ = 5). Kruskal-Wallis test with Dunn's multiple comparisons test; $p$ = 0.0179; $p$ > 0.9999 between E9.5 and E10.5; $p$ = 0.0761 between E9.5 and E11.5; $p$ = 0.0148 between E10.5 and E11.5.
**c)** Size of asymmetric neurogenic clones (number of cells) across induction time points. E9.5 ($n$ = 13), E10.5 ($n$ = 7), E11.5 ($n$ = 7). Kruskal-Wallis test with Dunn's multiple comparisons test; $p$ = 0.0750; p > 0.9999 between E9.5 and E10.5; $p$ = 0.1035 between E9.5 and E11.5; $p$ = 0.1732 between E10.5 and E11.5.
Violin plot data are presented as mean ± SEM. All clones were analyzed at E18.5. $n$ indicates the number of clones. *$p < 0.05$.

**Abbreviations:** SEM, standard error of the mean; E, embryonic day.

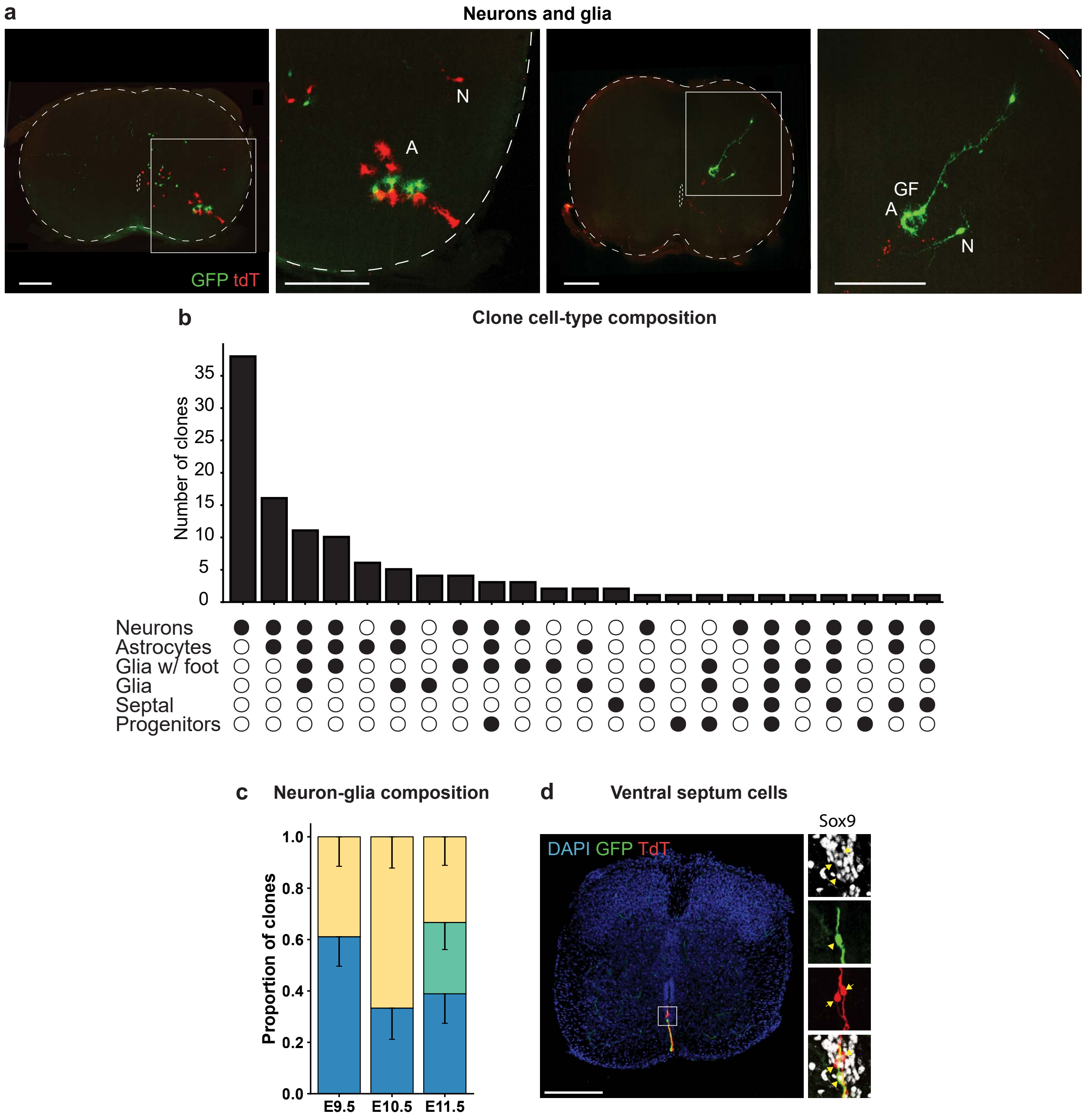
a
Neurons and glia
N
A
GF
A
N
GFP tdT
b
Clone cell-type composition
Number of clones
35
30
25
20
15
10
5
0
Neurons
Astrocytes
Glia w/ foot
Glia
Septal
Progenitors
c
Neuron-glia composition
Proportion of clones
1.0
0.8
0.6
0.4
0.2
0.0
E9.5
E10.5
E11.5
d
Ventral septum cells
DAPI GFP TdT
Sox9

**Supplementary Figure 3. MADM clone cell-type composition.**
**a)** Maximum intensity projections of MADM clones containing multiple cell types. Left: TM E11.5, cervical level; 250 μm projection; Right: TM E11.5, lumbar level; 150 μm projection. N: neuron; A: astrocyte; GF: glia with endfoot. tdT and GFP hemilineages are labeled
**b)** Frequency of cell-type compositions across all clones. Cells of uncertain identity excluded from quantification. *n* = 145.
**c)** Proportions of clones containing neurons and glia, neurons only or glia only across induction time points. Only clones with complete cell-type identification based on morphology are included. Data are presented as proportions with SE of the proportion (lower limit). E9.5 (*n* = 18), E10.5 (*n* = 15), E11.5 (*n* = 18). Fisher's exact test; $p = 0.1663$ between E9.5 and E10.5; $p = 0.0589$ between E10.5 and E11.5; $p = 0.0718$ between E9.5 and E11.5.
**d)** Spinning disk confocal image of cells in the ventral septum expressing Sox9. Scale bar, 200 μm Proportions are presented with SE of the proportion (lower limit). All clones were analyzed at E18.5. *n* indicates the number of clones.

**Abbreviations:** TM, tamoxifen; SE, standard error; E, embryonic day; GFP, green fluorescent protein; tdT, tdTomato; MADM, mosaic analysis with double markers; N + G, neurons and glia; G, glia-only; N, neurons-only.

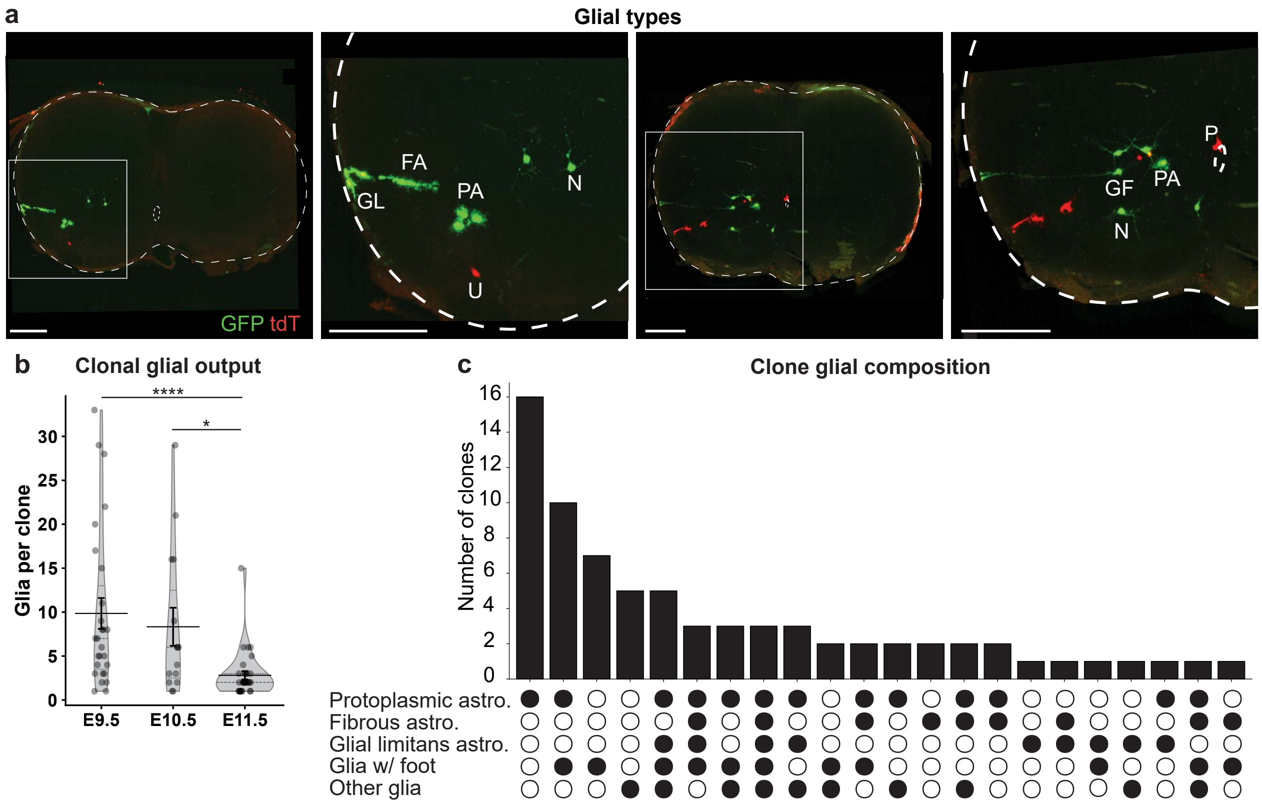

a
Glial types
GFP tdT
GL
FA
PA
N
U
P
GF
b
Clonal glial output
Glia per clone
****
*
E9.5
E10.5
E11.5
c
Clone glial composition
Number of clones
Protoplasmic astro.
Fibrous astro.
Glial limitans astro.
Glia w/ foot
Other glia

**Supplementary Figure 4. MADM clone glial composition.**
**a)** Maximum intensity projections of MADM clones containing multiple cell types. Left: TM E9.5, cervical level; 180 µm projection; Right: TM E9.5, cervical level; 150 µm projection.
**b)** Clonal glial output (number of glia per glia-containing clone) across induction time points. E9.5 (*n* = 27), E10.5 (*n* = 15), E11.5 (*n* = 32). Kruskal-Wallis test with Dunn's multiple comparisons test; $p < 0.0001$; $p > 0.9999$ between E9.5 and E10.5; $p < 0.0001$ between E9.5 and E11.5; $p = 0.0153$ between E10.5 and E11.5. □$p < 0.05$; □□$p$□ $< 0.0001$.
**c)** Frequency of glial compositions in glia-containing clones. Cells of uncertain identity excluded from quantification. *n* = 74.
Violin plot data are presented as mean ± SEM. All clones were analyzed at E18.5. *n* indicates the number of clones.

**Abbreviations:** TM, tamoxifen; SEM, standard error of the mean; E, embryonic day; GFP, green fluorescent protein; tdT, tdTomato; MADM, mosaic analysis with double markers; astro, astrocytes; N, neuron; PA, protoplasmic astrocyte; FA, fibrous astrocyte; GL, glial limitans astrocyte; U, uncertain morphology; WF, glia with endfoot; P, progenitor cell.

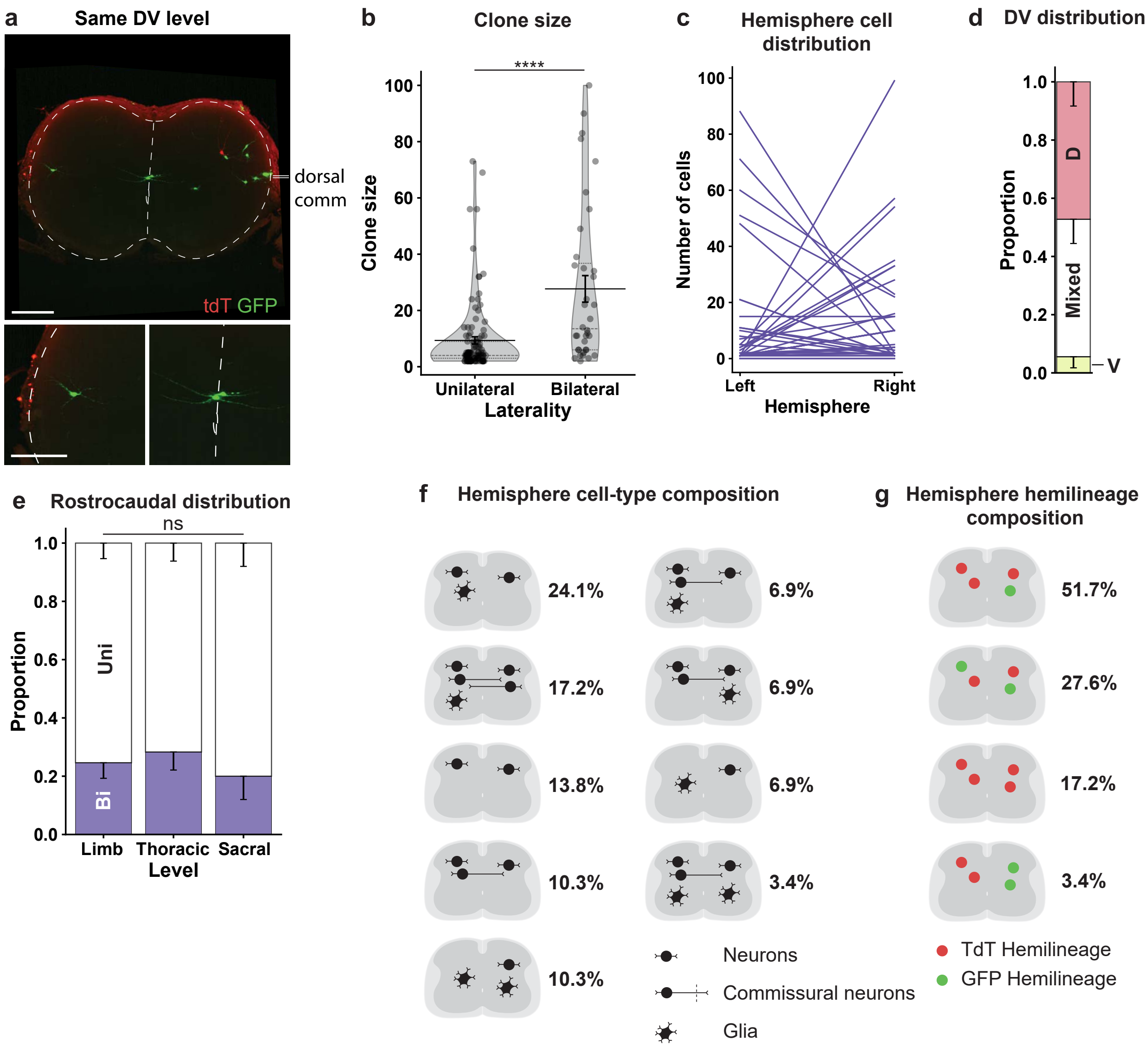

a Same DV level
dorsal comm
tdT GFP
b Clone size
****
Clone size
Unilateral Bilateral
Laterality
c Hemisphere cell distribution
Number of cells
Left Right
Hemisphere
d DV distribution
Proportion
D
Mixed
V
e Rostrocaudal distribution
ns
Proportion
Uni
Bi
Limb Thoracic Sacral
Level
f Hemisphere cell-type composition
24.1%
17.2%
13.8%
10.3%
10.3%
6.9%
6.9%
6.9%
3.4%
Neurons
Commissural neurons
Glia
g Hemisphere hemilineage composition
51.7%
27.6%
17.2%
3.4%
TdT Hemilineage
GFP Hemilineage

**Supplementary Figure 5. Bilateral clone position, size, and composition.**

**a)** Maximum intensity projection of a bilateral clone with contralateral neurons at a similar dorsoventral level. Cervical level; overview 350 µm projection; insets 150 µm projection. tdT and GFP hemilineages are labeled. The level of the dorsal commissure is indicated by double lines. Of note, this clone is the sole clone in a single-clone spinal cord.

**b)** Quantification of clone size (number of cells) in unilateral and bilateral clones. Unilateral (*n* = 107), bilateral (*n* = 36). Data are presented as means ± SEM. Mann-Whitney test; $p < 0.0001$.

**c)** Distribution of cells across the two hemispheres in bilateral clones. Each line represents the hemisphere distribution of one clone.

**d)** Proportion of bilateral clones situated dorsal to the central canal, ventral to the central canal, or spread across the dorsoventral axis.

**e)** Proportions of clones that are unilateral and bilateral at different rostrocaudal levels of the spinal cord. Data are presented as proportions with SE of the proportion (lower limit). Limb (*n* = 65), thoracic (*n* = 53), sacral (*n* = 25). Fisher's exact test; $p = 0.6788$ between limb and thoracic; $p = 0.5808$ between thoracic and sacral; $p = 0.7839$ between limb and sacral.

**f)** Percentage of hemisphere cell-type compositions in bilateral clones. Bilateral clones in which one hemisphere contains only cells without an identity assigned based on morphology are excluded from cell-type quantifications.

**g)** Percentage of hemisphere hemilineage compositions in bilateral clones.

Violin plot data are presented as mean ± SEM. Proportions are presented with SE of the proportion (lower limit). All clones were analyzed at E18.5. *n* indicates the number of clones. ****$p < 0.0001$.

**Abbreviations:** TM, tamoxifen; SE, standard error; SEM, standard error of the mean; E, embryonic day; GFP, green fluorescent protein; tdT, tdTomato; MADM, mosaic analysis with double markers; Bi, bilateral; Uni, unilateral; V, ventral; D, dorsal; DV, dorsoventral.

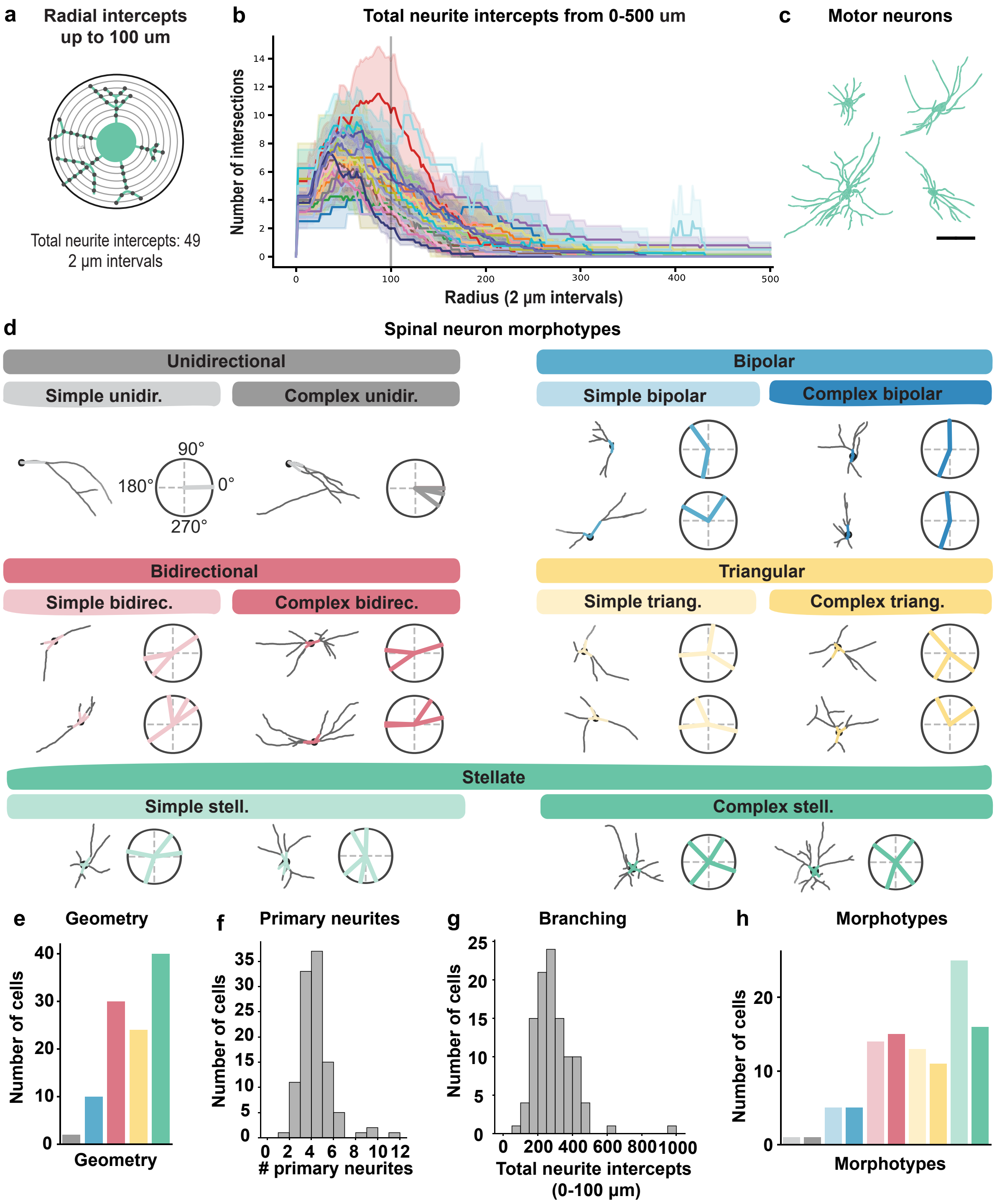

a
Radial intercepts up to 100 um
Total neurite intercepts: 49
2 µm intervals
b
Total neurite intercepts from 0-500 um
Number of intersections
Radius (2 µm intervals)
c
Motor neurons
d
Spinal neuron morphotypes
Unidirectional
Simple unidir.
Complex unidir.
90°
180°
0°
270°
Bipolar
Simple bipolar
Complex bipolar
Bidirectional
Simple bidirec.
Complex bidirec.
Triangular
Simple triang.
Complex triang.
Stellate
Simple stell.
Complex stell.
e
Geometry
Number of cells
Geometry
f
Primary neurites
Number of cells
### primary neurites
g
Branching
Number of cells
Total neurite intercepts (0-100 µm)
h
Morphotypes
Number of cells
Morphotypes

**Supplementary Figure 6. Defining spinal neuron morphotypes.**
**a)** Radial intercept analysis from 0–100 µm used to determine the total number of intercepts and assess neurite complexity.
**b)** Total neurite intercepts at 2 µm intervals from 0–500 µm across all reconstructed neurons (*n* = 106 neurons) from clones induced at E9.5 (*n* = 21) and E10.5 (*n* = 2). Most neurons peaked before 100 µm.
**c)** Motor neurons exhibit a complex stellate morphotype. Scale bar, 100 µm.
**d)** Example traces and neurite angle measurements for each morphotype.
**e–g)** Frequency distributions of neurite geometry (**e**), primary neurite number (**f**), and radial intercept counts (0–100 µm; **g**). Colour-code in **e** corresponds to titles in **d**.
**h)** Frequency of morphotypes across all reconstructed neurons. Colour-code corresponds to subtitles in **d**.
All clones were analyzed at E18.5. *n* indicates the number of clones, unless otherwise specified.

**Abbreviations:** TM, tamoxifen; E, embryonic day; unidir, unidirectional; bidirec, bidirectional; triang, triangular; stell, stellate.

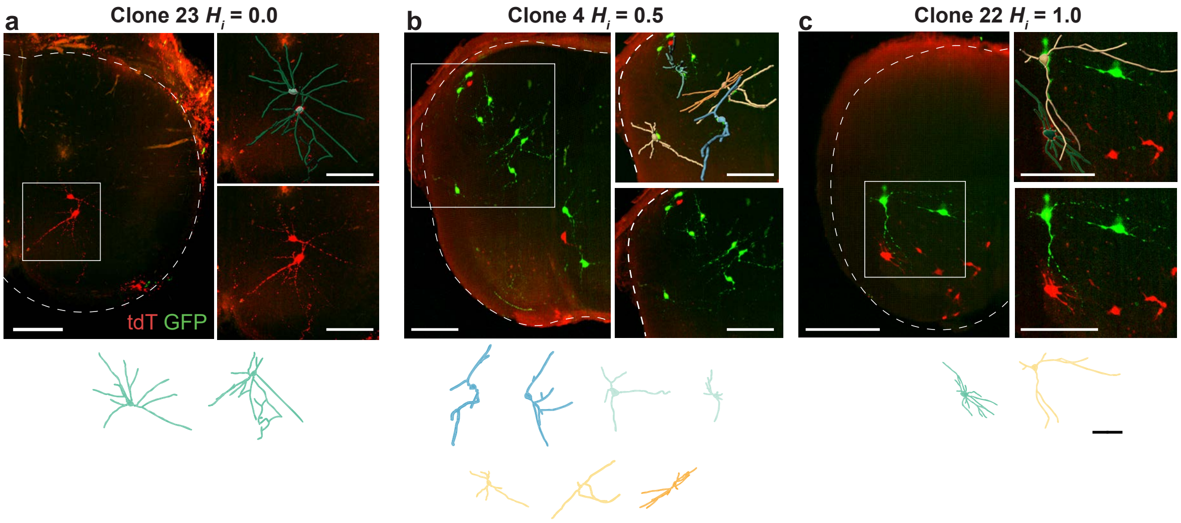
a
Clone 23 $H_i$ = 0.0
tdT GFP
b
Clone 4 $H_i$ = 0.5
c
Clone 22 $H_i$ = 1.0

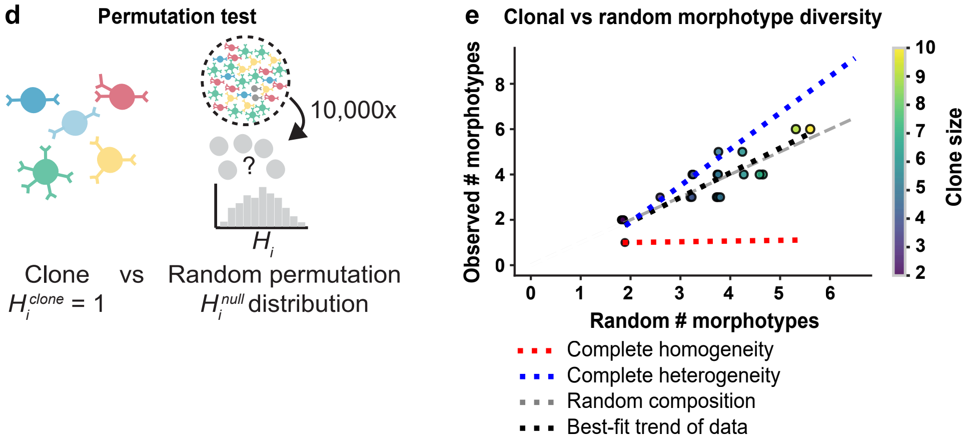
d
Permutation test
10,000x
?
$H_i$
Clone vs Random permutation
$H_i^{clone}$ = 1
$H_i^{null}$ distribution
e
Clonal vs random morphotype diversity
Observed # morphotypes
Random # morphotypes
Clone size
Complete homogeneity
Complete heterogeneity
Random composition
Best-fit trend of data

**Supplementary Figure 7. Morphotype heterogeneity in MADM clones.**

**a-c)** Representative MADM clones induced with TM at E9.5 and E10.5 with low (**a,** Cervical level; projection 400 μm), intermediate (**b,** Thoracic level; projection 500 μm) and high (**c,** Thoracic level; projection 300 μm) heterogeneity index, $H_i$, containing multiple morphotypes. Scale bars, 200 μm (image), 100 μm (traces). tdT and GFP hemilineages are labeled. Neuron trace colour-code corresponds to subtitles in **Fig. S6d**.

**d)** Schematic of the permutation test used to assess whether observed clonal heterogeneity differs from random expectation.

**e)** Relationship between observed morphotype number per clone and expected values from random permutations. For comparison, plot lines for hypothetical clones with complete homogeneity ($H_i$ = 0), complete heterogeneity ($H_i$ = 1), and random composition, are included. Colour bar indicates the number of reconstructed neurons per clone. A line of best fit closely matches a random composition. $r$ = 0.89.

All clones were analyzed at E18.5.

**Abbreviations:** TM, tamoxifen; E, embryonic day; GFP, green fluorescent protein; tdT, tdTomato; MADM, mosaic analysis with double markers.

**a**

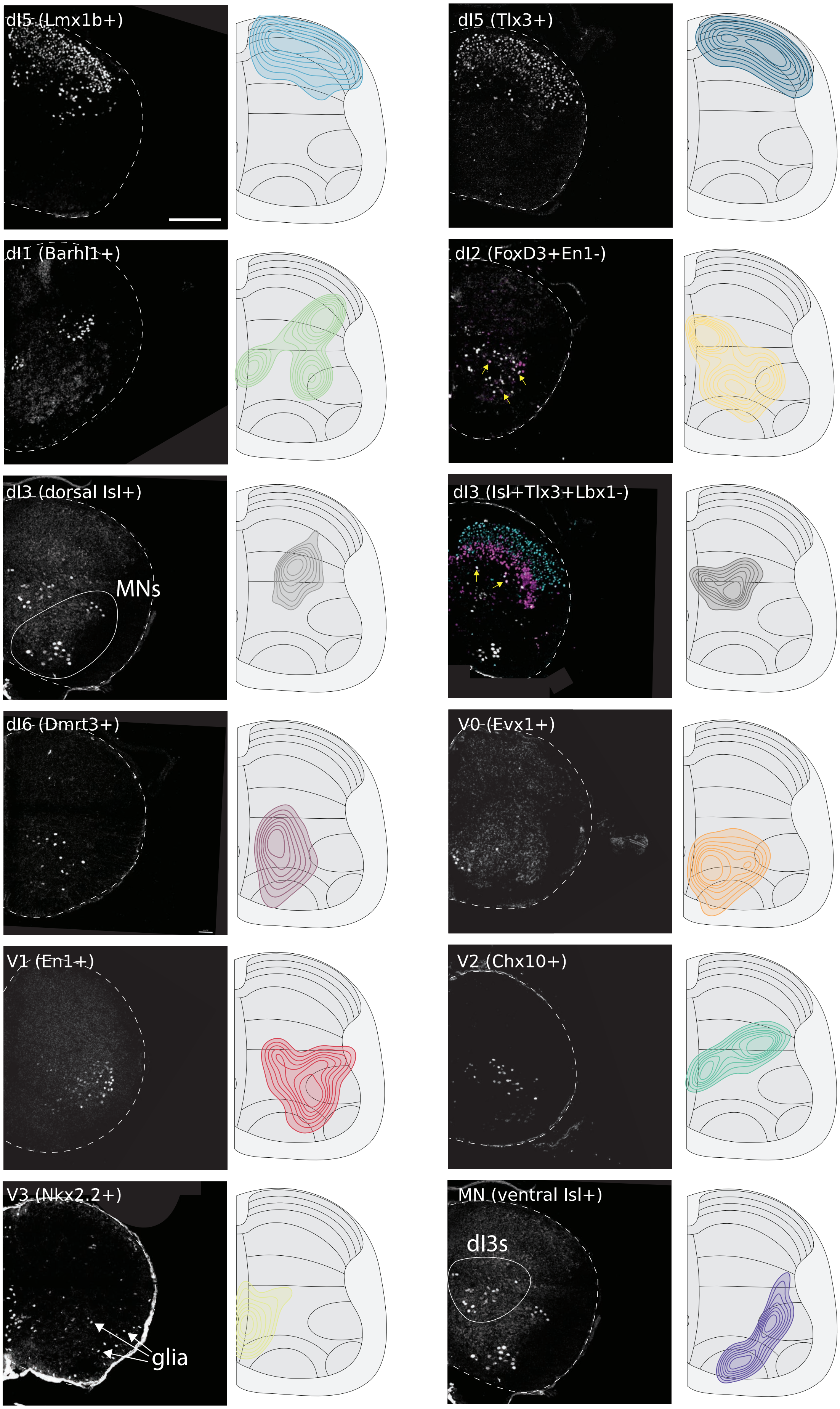

dI5 (Lmx1b+)
dI5 (Tlx3+)
dI1 (Barhl1+)
dI2 (FoxD3+En1-)
dI3 (dorsal Isl+)
MNs
dI3 (Isl+Tlx3+Lbx1-)
dI6 (Dmrt3+)
V0 (Evx1+)
V1 (En1+)
V2 (Chx10+)
V3 (Nkx2.2+)
glia
MN (ventral Isl+)
dI3s

**Supplementary Figure 8. Cardinal class spatial distribution map.**

**a)** Kernel density estimate (KDE) maps of cardinal classes at E18.5 based on immunofluorescence labeling of key transcription factor markers. The map captures Nkx2.2+ V3 interneurons but does not include the deep dorsal horn V3 subset (laminae IV–VI).

For each antibody, between 4 and 22 hemisections from between 1 and 4 animals were quantified.

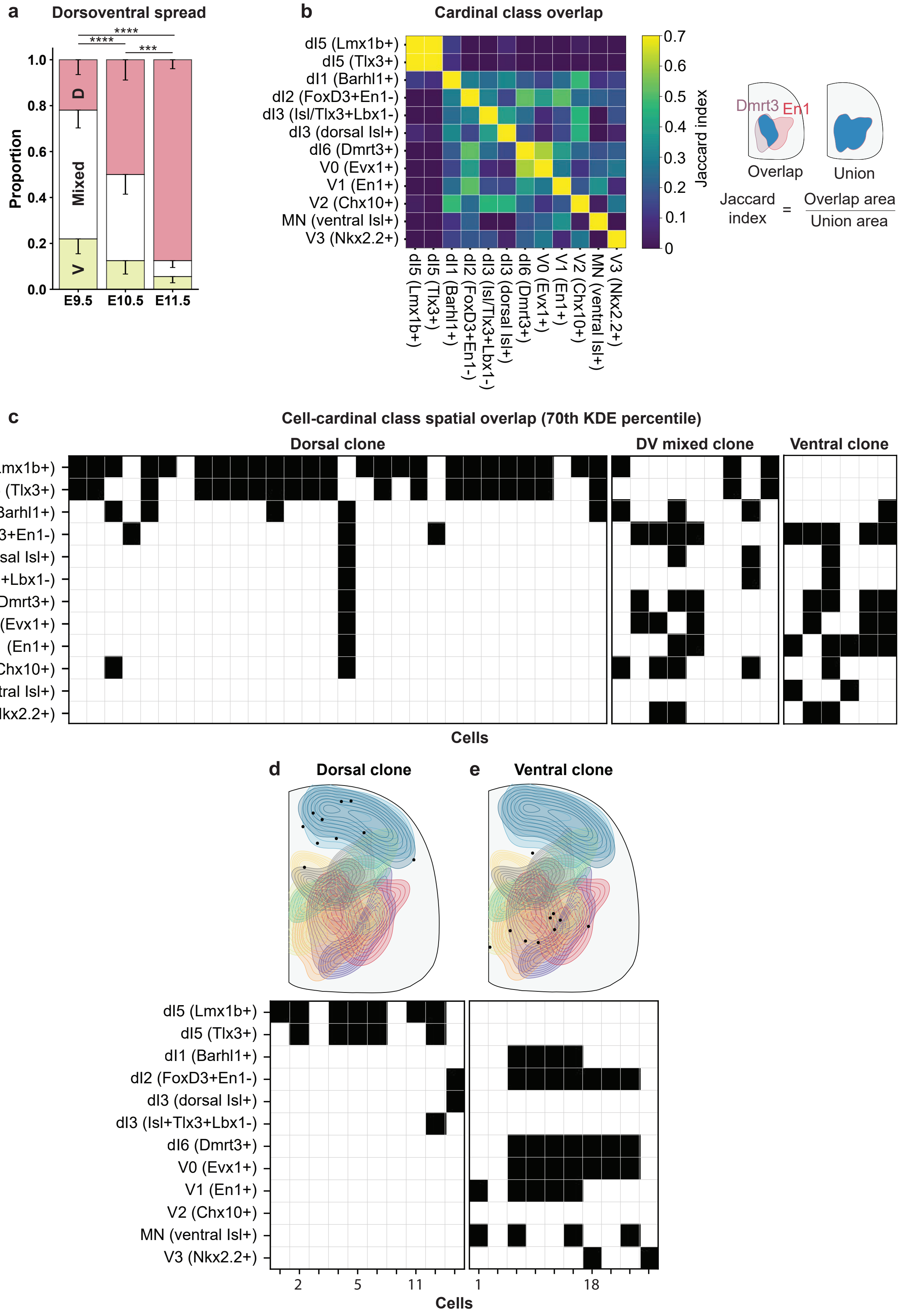

a
Dorsoventral spread
****
****
***
Proportion
1.0
0.8
0.6
0.4
0.2
0.0
D
Mixed
V
E9.5
E10.5
E11.5
b
Cardinal class overlap
dI5 (Lmx1b+)
dI5 (Tlx3+)
dI1 (Barhl1+)
dI2 (FoxD3+En1-)
dI3 (Isl/Tlx3+Lbx1-)
dI3 (dorsal Isl+)
dI6 (Dmrt3+)
V0 (Evx1+)
V1 (En1+)
V2 (Chx10+)
MN (ventral Isl+)
V3 (Nkx2.2+)
Jaccard index
0.7
0.6
0.5
0.4
0.3
0.2
0.1
0
Dmrt3
En1
Overlap
Union
Jaccard index = Overlap area / Union area
c
Cell-cardinal class spatial overlap (70th KDE percentile)
Dorsal clone
DV mixed clone
Ventral clone
dI5 (Lmx1b+)
dI5 (Tlx3+)
dI1 (Barhl1+)
dI2 (FoxD3+En1-)
dI3 (dorsal Isl+)
dI3 (Isl+Tlx3+Lbx1-)
dI6 (Dmrt3+)
V0 (Evx1+)
V1 (En1+)
V2 (Chx10+)
MN (ventral Isl+)
V3 (Nkx2.2+)
Cells
d
Dorsal clone
e
Ventral clone
dI5 (Lmx1b+)
dI5 (Tlx3+)
dI1 (Barhl1+)
dI2 (FoxD3+En1-)
dI3 (dorsal Isl+)
dI3 (Isl+Tlx3+Lbx1-)
dI6 (Dmrt3+)
V0 (Evx1+)
V1 (En1+)
V2 (Chx10+)
MN (ventral Isl+)
V3 (Nkx2.2+)
2
5
11
1
18
Cells

**Supplementary Figure 9. Class-class spatial overlap and predicted cell-class membership.**
**a)** Proportion of clones situated dorsal to the central canal (D), ventral to the central canal (V), or spread across the dorsoventral axis (Mixed) across labeling time points. E9.5 ($n$ = 41 clones), E10.5 ($n$ = 32), E11.5 ($n$ = 72). Fisher's exact test; $p < 0.0001$ between E9.5 and E10.5; $p$ = 0.0003 between E10.5 and E11.5; $p < 0.0001$ between E9.5 and E11.5.
**b)** Heatmap indicating the spatial overlap of cardinal class marker 70th kernel density estimate (KDE) percentile using the Jaccard index. Warmer colours indicate greater similarity in distributions.
**c)** Per-neuron heatmaps for the clones in **Fig. 4c,e** indicating the overlap of cells with the 70th KDE percentile of each class marker distribution.
**d-e)** Per-neuron heatmaps for the clones in **Fig. 4a-b** indicating the overlap of cells with the 70th KDE percentile of each class marker distribution.
Proportions are presented with SE of the proportion (lower limit). All clones were analyzed at E18.5. $n$ indicates the number of clones. ***$p < 0.001$; ****$p < 0.0001$.

**Abbreviations:** TM, tamoxifen; SE, standard error; E, embryonic day; GFP, green fluorescent protein; tdT, tdTomato; MADM, mosaic analysis with double markers; D, dorsal; V, ventral; KDE, kernel density estimate.
